# Locating cellular contents during cryoFIB milling by cellular secondary-electron imaging

**DOI:** 10.1101/2022.08.18.504468

**Authors:** Chao Lin, Li Zhang, Ziying Zhang, Yifeng Jiang, Xueming Li

## Abstract

Cryo-electron tomography (cryoET) is a powerful technique that enables the direct study of the molecular structure of tissues and cells. Cryo-focused ion beam (cryoFIB) milling plays an important role in preparation of high-quality thin lamellar samples for cryoET studies, promoting the rapid development of cryoET in recent years. However, locating the regions of interest in a large cell or tissue during cryoFIB milling remains a major challenge limiting cryoET applications on arbitrary biological samples. Here, we report an on-the-fly location method based on cellular secondary electron imaging (CSEI). CSEI is derived from a basic imaging function of the cryoFIB instruments and enables high-contrast imaging of the cellular contents of frozen hydrated biological samples, highlighted by that both fluorescent labels and additional devices are not required. The present work discusses the imaging principles and settings for optimizing CSEI. Tests on several commercially available cryoFIB instruments demonstrated that CSEI was feasible on mainstream instruments to observe all types of cellular contents and was reliable under different milling conditions. Assisted by CSEI, we established a simple milling-location workflow and tested it using the basal body of *Chlamydomonas reinhardtii*.

## Introduction

Cryo-electron tomography (cryoET) is a popular technique that allows the observation of the *in situ* molecular structure of tissues and cells. However, owing to the weak penetration of electrons, cryoET works only on thin samples, typically 100–200 nm in thickness. Meanwhile, the view field of a single cryoET snapshot is usually smaller than 1 μm, which is limited by the available number of pixels on the camera and the desired resolution. Therefore, the total volume of a cryoET tomogram is typically less than 0.1 μm^3^. The development of cryo-focused ion beam (cryoFIB) milling has largely solved the problem of preparing such a small and thin lamella, thereby promoting the rapid development of cryoET in recent years. However, locating such a tiny region inside frozen tissues or cells with a size of thousands of cubic micrometers is still challenging.

The currently available methods for locating the regions of interest in cryoET sample preparation are mainly based on fluorescence, known as cryo-correlated light and electron microscopy (cryoCLEM)**^1–4^**. CryoCLEM matches the images of fluorescence light microscopy and cryo-electron microscopy (cryoEM) and then locates the regions of interest in an electron microscope based on fluorescence light microscopy images. CryoCLEM can assist cryoFIB milling in two ways, i.e., by using separated optical instruments or by integrating optical microscopy attachments into a cryoFIB instrument. The former does not allow on-the-fly positioning during milling. The resolution of fluorescence imaging is much lower than that in conventional applications because of the long working distance of the optical objective lens, which is required for liquid nitrogen cooling and to avoid contamination. In the second method, on-the-fly positioning is possible using an integrated optical microscope, but the optical resolution is much lower than that of the first approach because of the similar working distance issue and limited space inside the cryoFIB instrument. While both confocal and super-resolution fluorescence microscopy have been used in cryoCLEM and have the potential to achieve three-dimensional (3D) localization, localization in the axial direction is still challenging, mostly limited by poor resolution**^5, 6^**. In addition to these problems, the complicated procedure of cryoCLEM operations and the lack of stable commercial devices are major limiting factors in practice. However, devitrification due to optical excitation**^5–7^** and the need for fluorescence labels are sometimes issues for some samples.

Therefore, finding an alternative method, especially enabling on-the-fly location during cryoFIB milling, is necessary and important. Secondary electron imaging may be an ideal candidate. CryoFIB instruments are usually designed based on scanning electron microscope that simultaneously supports scanning electron microscopy (SEM) imaging. The secondary electrons are the most basic signals used in SEM imaging and usually provide topographical imaging to assist in cryoFIB milling. Secondary electron excitation is insensitive to element composition and is seldom used for composition-related imaging. Some studies on cryoFIB-SEM block face imaging**^8–12^** reported secondary electron imaging of frozen hydrated biological samples, which showed high contrast of the cellular contents, including organelles and membranes, on flat surfaces prepared by cryoFIB milling. The contrast of secondary electron images is thought to be related to the water content and lipid composition, thus exhibiting the ultrastructure of the cells. The mechanism of contrast formation is complicated and related to the interactions between the primary electrons and the exposed biological sample after milling**^8^**. Because secondary electron imaging is the most fundamental function of a cryoFIB instrument, it would be an ideal solution for on-the-fly 3D location.

Herein, we report a method based on cellular secondary electron imaging (CSEI) for accurate on-the-fly 3D location of frozen hydrated biological samples during cryoFIB milling. This method does not require fluorescent markers, special sample processing, or additional devices. We established a complete workflow for CSEI-based location, compared and optimized the imaging quality of several commercially available cryoFIB instruments. Samples from different species, including bacteria, *Chlamydomonas,* mammalian cells, mammalian and plant tissues, were tested to demonstrate CSEI use in locating organelles, membraneless organelles, and protein aggregates. Finally, we demonstrate a complete 3D locating-milling workflow using the basal body of *C. reinhardtii* flagella.

## Results

### Secondary electron imaging for frozen hydrated cellular samples

SEM uses primary electron beam scanning across the sample surface to excite detectable signals, such as secondary and backscattered electrons, for imaging. The excited secondary electrons are the signals used in the present work and can be classified into at least three types**^13^**: those excited on the sample surface directly by the primary electron beam (termed SE_I_); those excited by the backscattered electrons inside the sample (termed SE_II_); and those excited by the backscattered or primary electrons striking the chamber or polepiece (termed SE_III_ and SE_IV_). SE_I_ and SE_II_ are generated from the sample and play a major role in imaging. Secondary electrons have low energy (typically less than 50 eV) and can only escape from the shallow surface (typically less than 10 nm) of the sample**^13^**. In topographical imaging applications during cryoFIB milling, secondary electron images are mostly optimized to display the shape of the sample after milling rather than to observe the cellular structures in the milled surface.

In several reports on cryoFIB-SEM block face imaging, cellular contrast by the secondary electron imaging has been observed on cryoFIB-milled surfaces. Cellular contrast originates from the escaping secondary electrons on the sample surface. The more electrons that escape, the brighter the corresponding area is. Many factors can influence electron escape or contrast formation, including but not limited to the follows**^8^**: (a) a negatively charged surface promotes secondary electron escape; conversely, a positively charged surface suppresses emission; (b) the electric state inside the biological sample, such as hydrophilic and hydrophobic interactions, also affects the efficiency of secondary electron production**^8^**; (c) the production efficiency of the secondary electrons is relatively sensitive to the element type for light atoms (atomic number less than 20); hence, the biological sample might exhibit some compositional contrast. Combining these properties of secondary electron escape, we refer to the related imaging formation as CSEI and use CSEI to observe and locate different cellular contents (**Fig. 1**). Regions with high water content, such as the interstitial spaces of intracellular materials, were shown in bright grayscale. Vesicles with relatively higher water content (**Fig. 1a**) showed higher brightness. In contrast, membrane structures (**Fig. 1b**), organelles with dense proteins (**Fig. 1c**), and protein condensates, such as starch sheaths (**Fig. 1d**) and chromatin aggregates (**Fig. 1e**), were displayed in black grayscale. These high-contrast features enabled the location of cellular contents.

**Figure 1.**
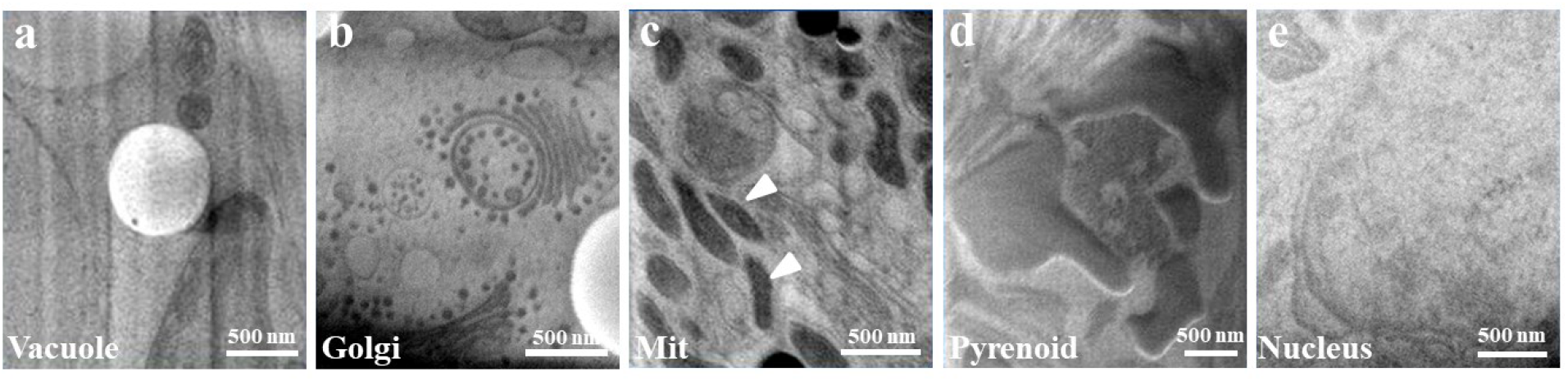
CSEI of different cellular contents. **a**, A vesicle in *C. reinhardtii* cell. **b**, Golgi apparatus in *C. reinhardtii* cell. **c**, Mitochondria in HeLa cell, pointed by white arrows. **d**, A pyrenoid in *C. reinhardtii* cell. **e**, Nucleus in HeLa cell. All images were collected using Crossbeam 550.

### Hardware configurations for imaging frozen hydrated samples

Secondary electron imaging is usually optimized for topographical imaging to assist cryoFIB milling (**Supplementary Fig. 1a-c**) and, hence, seldom shows cellular contrast with the default settings on some instruments (**Supplementary Fig. 1d**). The hardware configurations and imaging settings (discussed in the next section) should be considered to enable the location of the CSEI.

We tested several cryoFIB instruments for CSEI, including Helios (Thermo Fisher Scientific), Aquilos 1 and 2 (Thermo Fisher Scientific), and Crossbeam 550 (Carl Zeiss Microscopy GmbH). Multiple secondary electron detectors were installed at different locations inside these instruments and categorized into in-lens and in-chamber detectors (**Supplementary Table 1**). These detectors could only detect a portion of the secondary electrons that escaped in specific directions. The in-lens detectors were installed inside the lens or column and mainly detected electrons escaping at high angles (relative to the sample surface), mostly SE_I_. SE_II_ had a wide range of escape directions and were mainly detected by an in-chamber detector. SE_I_ were more sensitive to the surface electronic potential of the sample because the surface potential was perpendicular to the surface, that is, along the escape direction of the SE_I_. The resolution of SE_II_ imaging was generally lower than that of SE_I_ imaging because the backscattered electrons had a larger interaction area in the sample than the primary electrons. In summary, the images from the in-lens detectors often have better resolution than those from the in-chamber detectors but are more sensitive to surface charging**^13^**. However, the actual imaging efficacy is complicated. For example, the incident direction of the primary electrons is usually not perpendicular to the milled sample surface in all the tested cryoFiB instruments, which complicates the relationship between the detection and the escape angles of the secondary electrons. Morever, the detection principle, detection position, and parameter settings of the detectors on different instruments can vary, leading to significant differences in the imaging results (**Fig. 2**, **Supplementary Fig. 2**, and **Supplementary Fig. 3**).

**Figure 2.**
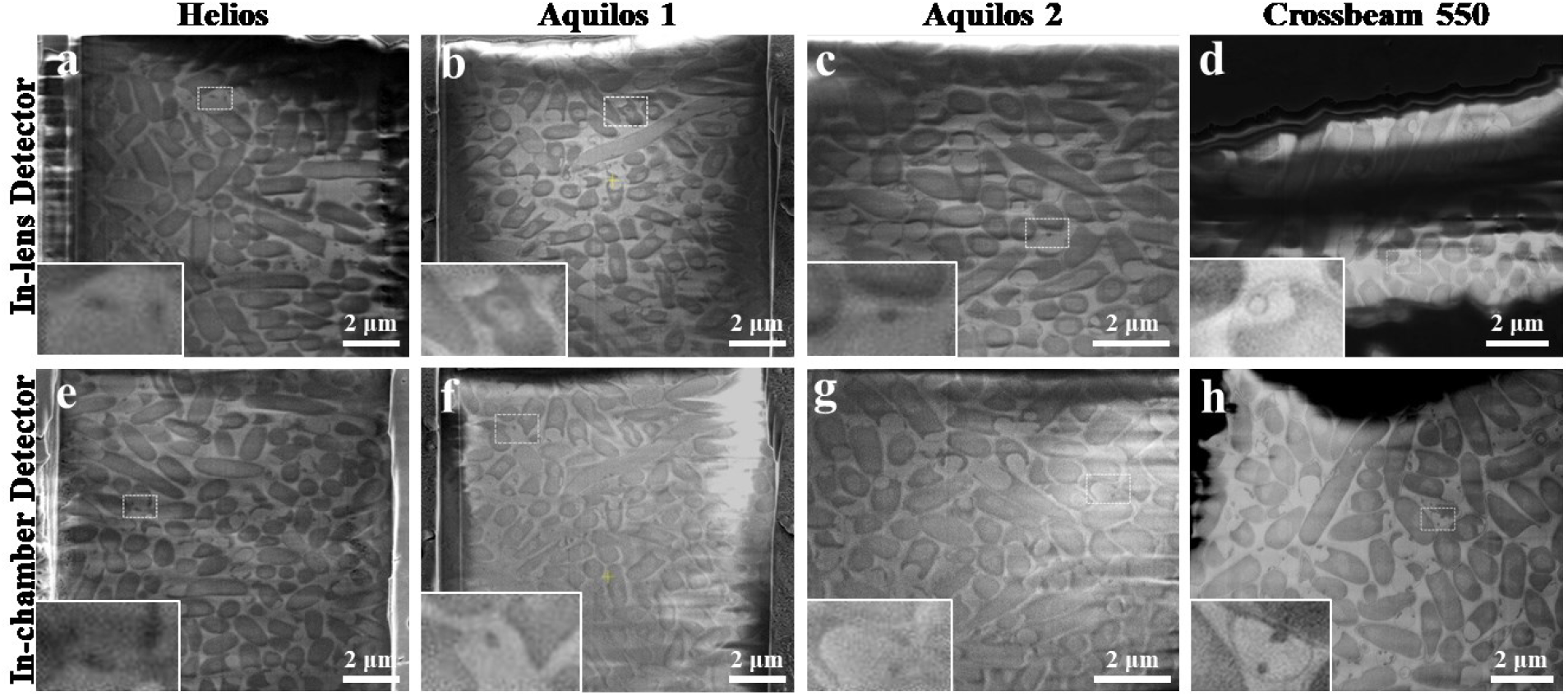
*E. coli* cells visualized by different detectors of several cryoFIB instruments. **a, b, c,** and **d,** Typical frozen *E. coli* images collected by the in-lens detectors of four cryoFIB instruments. **e**, **f**, **g**, and **h**, Typical frozen *E. coli* images collected by the in-chamber detectors of four cryoFIB instruments. The name of the corresponding cryoFIB instrument for each column is labeled on the top. In each image, a small rectangle region is magnified and inset in the bottom left, which shows fine features of the membrane and cell boundary. All the images were acquired with the optimized image settings and demonstrated the best imaging quality that we could obtain on corresponding instruments.

All cryoFIB instruments tested here have the capability of CSEI for frozen hydrated cellular samples. In the tests using *Escherichia coli* samples (**Fig. 2**), Crossbeam 550 showed the best resolution (**Fig. 2d, h**). As expected, the in-lens detectors (**Fig. 2a-d**) often have better resolution than the in-chamber detectors (**Fig. 2e-h**) but are more frequently influenced by shadows (**Fig. 2a-d**) associated with surface charging. Some instruments support the simultaneous output of separated images from different detectors. We can either choose a single image or merge them to generate a more complete image to minimize the influence of shadows. For example, the shadow areas from the in-lens (**Supplementary Fig. 4a**) and in-chamber (**Supplementary Fig. 4b**) detectors of Crossbeam 550 are often complementary and can be dismissed by merging the images from the two detectors (**Supplementary Fig. 4c**).

In addition, different lens settings are often used for survey and high-resolution imaging mode (**Supplementary Table 2**). These settings include the lens mode, beam size, and the effective working distance of the lens. The survey mode usually aims to provide a fast and large view at a lower resolution than the high-resolution mode. In our test, the high-resolution mode presented clearer features of the outer membrane than did the survey mode (**Supplementary Fig. 5**). However, the resolution loss of the survey mode seems modest; hence, it should still be sufficient for most location purposes. A shorter working distance has a positive influence on the imaging resolution of the in-lens detectors but is quite subtle (**Supplementary Fig. 6-7**). We also observed that the high-resolution mode was often severely affected by surface charging compared with the survey mode when using the in-lens detector on Helios (**Supplementary Fig. 5c** and **Supplementary Fig. 6d, h, l**).

### Imaging settings for frozen hydrated samples

In addition to hardware configurations, the imaging settings also play an important role in displaying cellular features. It is necessary to optimize these settings to obtain high-quality CSEI.

First, the acceleration voltage of the primary electron beam is a key factor (**Supplementary Fig. 8-10**). Reducing the incident electron energy can increase secondary electron emission**^13^**. However, a lower voltage reduces the penetration capability of primary electrons, leading to a decrease in the electron interaction depth of the sample**^13^**. Consequently, the secondary electron signal excited by the primary electrons with a lower voltage becomes more sensitive to the extreme surface state of the sample**^13^**. We tested an accelerating voltage of 1 kV, all images had poor contrast (**Supplementary Fig. 8a-d**, **Supplementary Fig. 9a-c,** and **Supplementary Fig. 10a-b**), even showing carved features (**Supplementary Fig. 9a-c**). One explanation was that the secondary electron signal excited at such a low voltage was mainly from the shallow surface that was damaged by FIB radiation. When the voltage was increased to 2 or 3 kV, the interaction depth increased, allowing the signal of the undamaged biological structures under the damaged surface layer to be excited (**Supplementary Fig. 8e-l**, **Supplementary Fig. 9d-f,** and **Supplementary Fig. 10c-f**). Upon further increasing the voltage to 5 kV, the charging problem was obviously enhanced (**Supplementary Fig. 8m-p**, **Supplementary Fig. 9g-i,** and **Supplementary Fig. 10g-h**). This might indicate charging accumulation in the bulky sample because the number of electrons escaping from the sample became less than the number of input primary electrons at a high accelerating voltage (corresponding to the upper crossover energy E2 given by Joy and Joy**^14^**). In addition, the cellular contrast of some images taken at 1 kV was reversed relative to the images obtained at higher voltage (**Supplementary Fig. 8a-d**). This phenomenon might also be related to the balance between the amount of input and the escape of electrons**^13^**. In summary, an acceleration voltage of 2–3 kV was the choice for the CSEI.

Secondly, increasing the electron beam dwell time (**Supplementary Fig. 11-12**) and electron beam current (**Supplementary Fig. 13**), as well as increasing the number of repetitive scans (**Supplementary Fig. 14**), could improve the imaging contrast. This improvement should benefit from a better signal-to-noise ratio by inceasing the radiation dose. More radiation causes greater radiation damage and charge accumulation inside the sample body. A longer dwell time and repetitive scans increase the imaging time, hence, make the image susceptible to the sample motion (**Supplementary Fig. 11e, f** and **Supplementary Fig. 14i, j**), ultimately, affecting the imaging resolution. We typically use an electron beam current of 50 pA, a dwell time of 1 μs and repetitive scans of 20 times.

### Secondary electron imaging and 3D locating during cryoFIB milling

On-the-fly 3D location can be enabled by frequently applying a CSEI during FIB milling. By imaging each milled surface, we were able to search for target objects based on cellular features. Initially, a large ion beam current, typically as high as 3 nA, was applied on a large area to achieve fast initial milling and searching, followed by precise milling with progressively reduced ion beam currents (**Fig. 3a**) to improve sample surface flatness and decrease radiation damage, as well as to further pinpoint the target location. Such a milling procedure requires CSEI on various surfaces generated by different ion beam currents.

**Figure 3.**
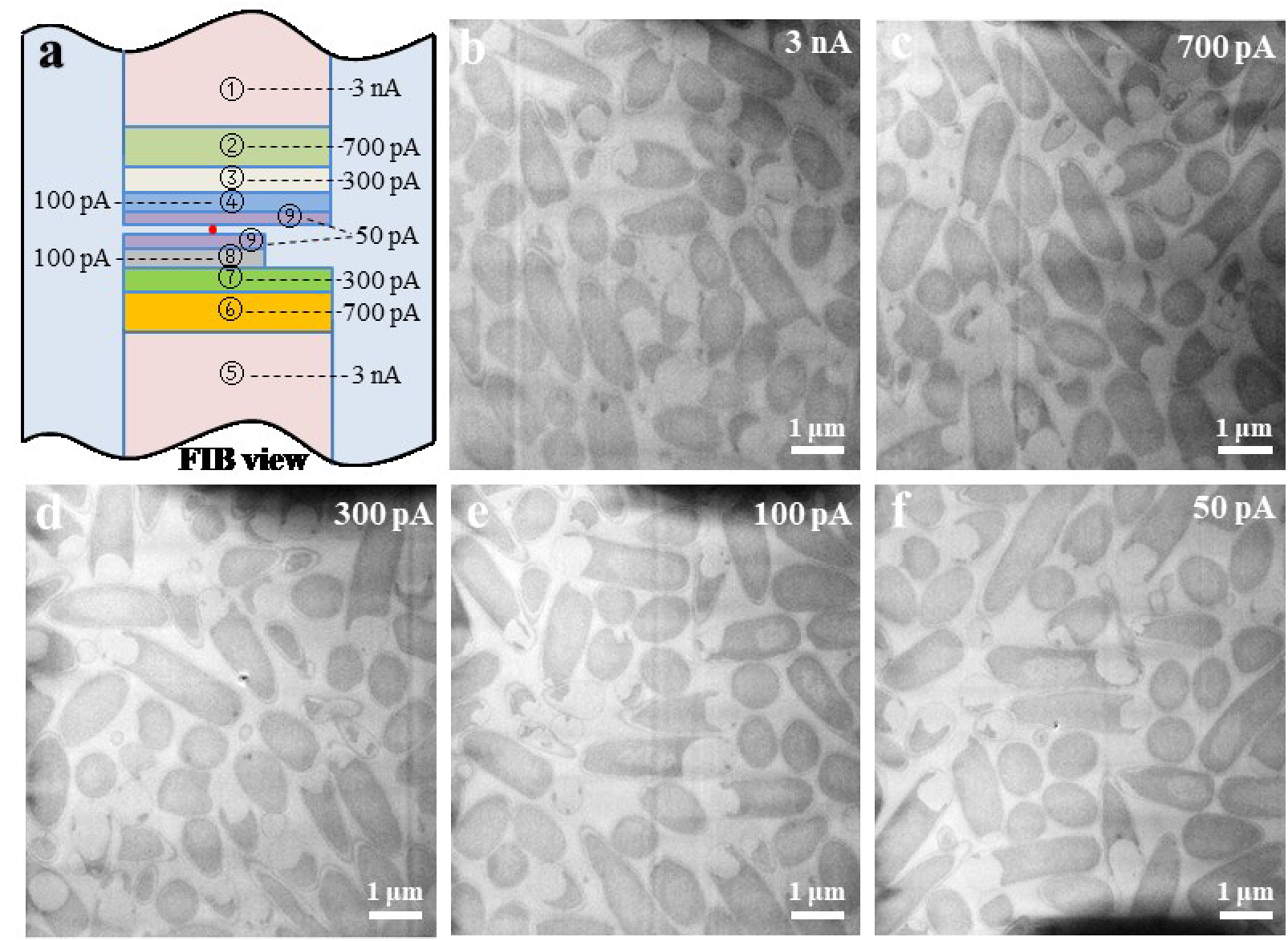
The milling-locating workflow and CSEI on the surface milled by different ion beam currents. **a**, The milling patterns are shown under the FIB view, illustrating the milling-locating workflow. The blue volume represents the remaining sample volume. Strips with other colors indicate the volume removed by cryoFIB milling. The associated numbers of strips indicate the milling sequence with the corresponding ion beam current. The red dot on the final lamella presents the object of interest. **b**, **c**, **d**, **e**, and **f**, CSEI on the surfaces of frozen hydrated *E. coli,* milled by different ion beam currents shown on the top right. All images were collected using Crossbeam 550.

A higher cryoFIB current may cause more surface radiation damage and hence influence the imaging quality. In addition, a high ion beam current often produces a rough surface and makes the milling sensitive to surface ice contamination, which is known as the ‘curtaining issue’**^15^**. We tested the CSEI with weak and strong curtaining issues under different ion beam currents. The variation of the ion beam currents and the presence of curtain did not influence the CSEI, as demonstrated by the precise cellular features of the bacteria (**Fig. 3b-f** and **Supplementary Fig. 15**).

The resolution of the CSEI was at the nanometer level, which was sufficient to resolve most membrane structures. Such resolution enables the visualization of tightly interacting membranes, such as the inner and outer membranes of *E. coli* (**Fig. 4a**), as well as the stacked thylakoids in the chloroplasts of *C. reinhardtii* (**Supplementary Fig. 16**).

**Figure 4.**
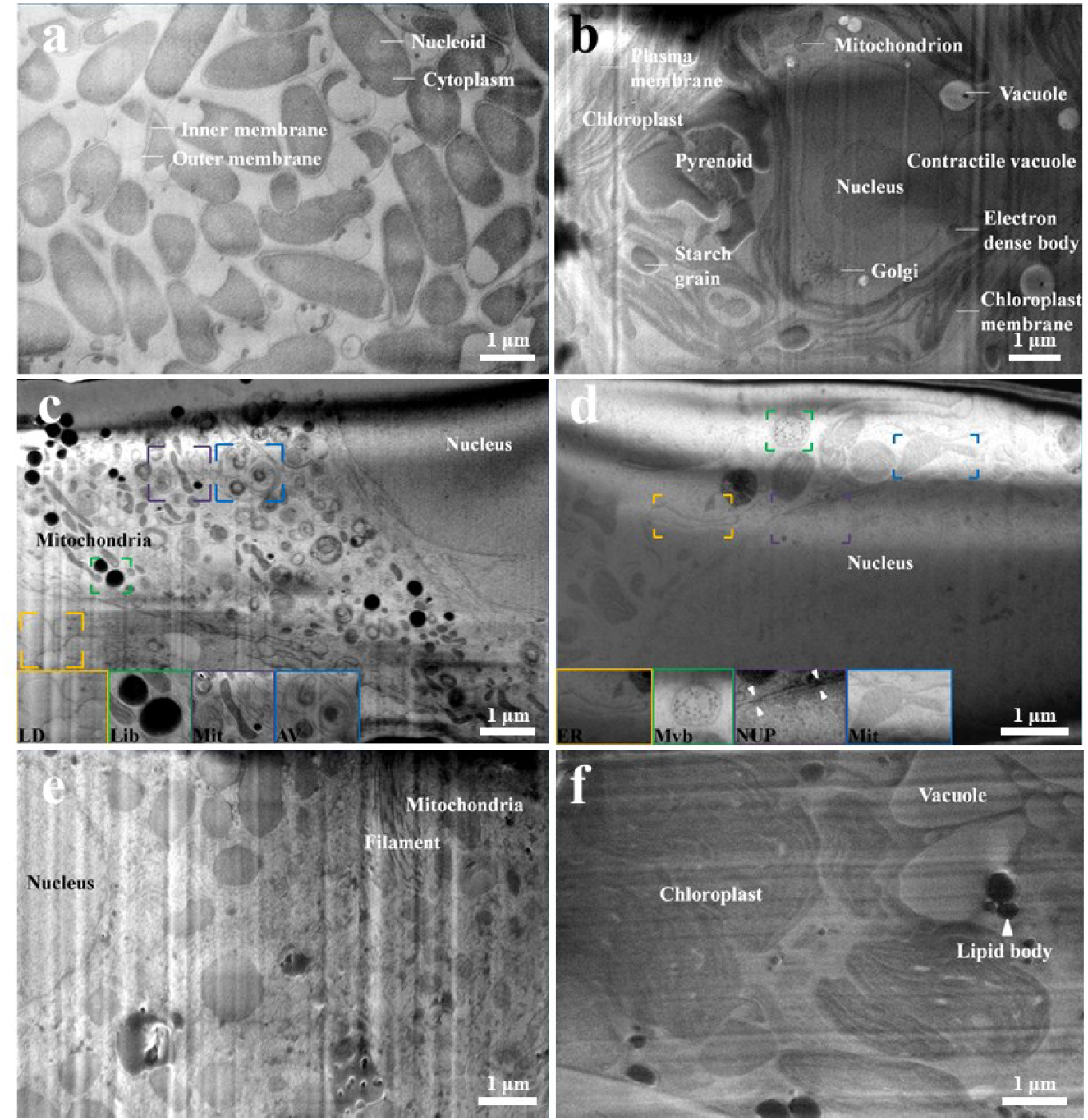
Various cellular contents visualized by CSEI. **a**, *E. coli* cells. **b**, A *C. reinhardtii* cell. **c**, A human skin squamous carcinoma cell. Insets show enlarged views of lipid droplets (LD), lipid body (lib), mitochondria (Mit), and autophagic vesicles (AV). **d**, A HeLa cell. Insets show enlarged views of endoplasmic reticulum (ER), multivesicular bodies (mvb), nucleopores (NUP), and mitochondria (Mit). **e**, A mouse liver cell. **f**, A plant cell in a *R. sativus* tissue. Recognized organelles are labeled with their names in the images. All images were collected using Crossbeam 550.

### Secondary electron imaging of organelles in single cells and tissues

To locate the intracellular organelles in the frozen state, we tested the CSEI using several different cell samples. In prokaryotic *E. coli*, the inner and outer membranes, vesicles, and cavities could be clearly distinguished, and the nucleoid had a brighter gray level than cytoplasm (**Fig. 4a**). In unicellular eukaryotic *C. reinhardtii* cells, characteristic features were observed, including a cup-shaped chloroplast occupying half of the cell, a pyrenoid located at the base of the chloroplast, and a nucleus surrounded by the chloroplast (**Fig. 4b**). Meanwhile, organelles such as Golgi bodies, Golgi vesicles, mitochondria, vesicles, vacuoles, contractile vacuoles, electron dense bodies, and starch grains could also be clearly identified (**Fig. 4b**). In mammalian human skin squamous carcinoma (A431 cells) (**Fig. 4c**) and HeLa (**Fig. 4d**) cells, the nucleus occupies a large volume of the cell, in which the double-layered nuclear membrane and even the nuclear pores could be clearly distinguished. Organelles such as mitochondria, lipid droplets, lipid bodies, multivesicular bodies, endoplasmic reticulum, and autophagic vacuoles could be clearly identified, and even ridges inside the mitochondria could be observed (**Fig. 4c, d**).

Bulk samples of plant and animal tissues were observed. The imaging quality of the tissue samples was not as good as that of the single-cell samples. The reason for this remained unclear. Nonetheless, various organelles were still clearly visible, including filaments in the mouse liver tissue (**Fig. 4e**). In *Raphanus sattvus* plant tissues, membrane structures within the chloroplasts were clearly visible (**Fig. 4f**). In another observation for the same sample, some chloroplasts were slightly lighter than others, which might be related to the different contents of the chloroplast matrix proteins (**Supplementary Fig. 17**).

These tests demonstrate that the CSEI can be used to observe and locate organelles and their fine structures and hence is generally applicable to samples from different species.

### Secondary electron imaging of membraneless organelles and aggregates

Studies on membraneless organelles and protein aggregates inside cells are popular in cell biology. CSEI provides a way to precisely locate these cellular contents. We first tested *C. reinhardtii* cells and observed the phase separation droplet formed by Rubisco interacting with EPYC1**^16^**. The droplet was wrapped by a pyrenoid and was clearly distinguished from other regions of the cells (**Fig. 5a**). In another *C. reinhardtii* cell, we observed that the Rubisco droplet contained some texture features and did not fill the entire pyrenoid interior, which was related to the different developmental stages of the cells**^17^**(**Fig. 5b**). The nucleolus is a membraneless cellular compartment that is thought to be associated with phase transitions**^18^**. We observed distinctly different contrasts in the nucleolus of *C. reinhardtii* and HeLa from the surrounding nucleoplasm by CSEI (**Fig. 5c-d**). In addition to the nucleolus, chromatin in different aggregation states, including heterochromatin and euchromatin, were observed with different grayscales (**Fig. 5e**). In *E. coli* overexpressing the *Thermoplasma acidophilum* 20S (T20S) proteasome, protein aggregates frequently appeared in dark contrast (**Fig. 5f, g**). These results suggest that CSEI can be used to observe and localize membraneless organelles and protein aggregates.

**Figure 5.**
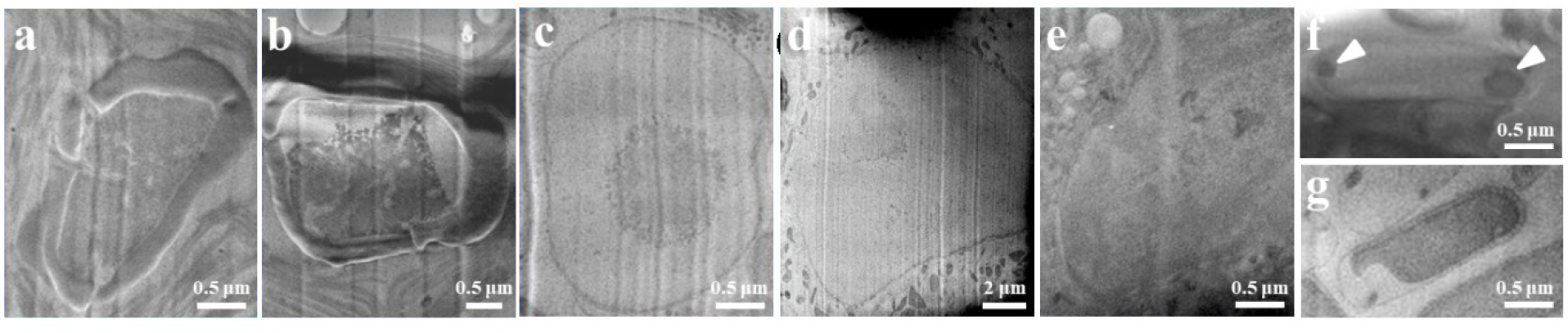
Membraneless organelles and protein aggregates visualized by CSEI. **a,** and **b,** Rubisco phase separation droplets at different developmental stages of *C. reinhardtii* cell. **c,** A nucleolus of *C. reinhardtii* cell. **d**, A nucleolus of a HeLa cell. **e**, Variable nuclear densities in a normal rat kidney (NRK) cell. **f**, A *E. coli* cell overexpressing T20S proteasome. Protein aggregations presented at the two ends of the bacterial cell are pointed by white arrows. **g**, A *E. coli* without overexpressed T20S proteasome. All images were collected using Crossbeam 550.

### An example of locating and precisely milling the basal body of *C. reinhardtii*

The *C. reinhardtii* basal body is the organizing center of the flagellum**^19^** and has a diameter of approximately 250 nm**^20^**. Mature *C. reinhardtii* contain only one basal body pair. The low amount and small size compared with the ~10 μm cell size make it nearly impossible to prepare a thin lamella containing the basal body without on-the-fly locating. We demonstrated a cryoFIB milling procedure with CSEI using the basal body as the target.

Cultured *C. reinhardtii* cells were plunge-frozen and Pt-coated, following a general protocol (see Methods). The initial cryoFIB milling was performed in a window of 20 μm width and 7 μm height under the FIB view (**Supplementary Fig. 18a**), using a large 3 nA ion beam current. The first CSEI showed clear structures in several cells (**Fig. 6a**). The basal body is typically found on the *C. reinhardtii* head that is characterized by the adjacent nucleus and vesicles. Based on these features, we determined a targeting position (**Fig. 6a**) and performed millings with a depth step of ~0.6 μm and an ion beam current of 700 pA. After repeating this milling process twice (**Fig. 6b-c**), we performed multiple milling processes with a smaller depth step of ~0.3 μm and a smaller ion beam current of 300 pA. After removal at a depth of more than 2.8 μm (relative to the surface of **Fig. 6a**), we observed the basal body (**Fig. 6d**). In such a milling-locating procedure, the choice of milling steps and ion beam currents are determined according to the actual situations. A smaller step and a lower current should be used closer to the predicted targeting depth. After reaching the target, we milled the opposite side of the sample and reduced the lamellar width to 12 μm to improve the milling efficiency (**Fig. 6e** and **Supplementary Fig. 18**). After finishing the milling on both sides of the lamella, the lamella is usually polished by further milling to a 20 nm depth on two surfaces with a small ion beam current of 50 pA.

**Figure 6.**
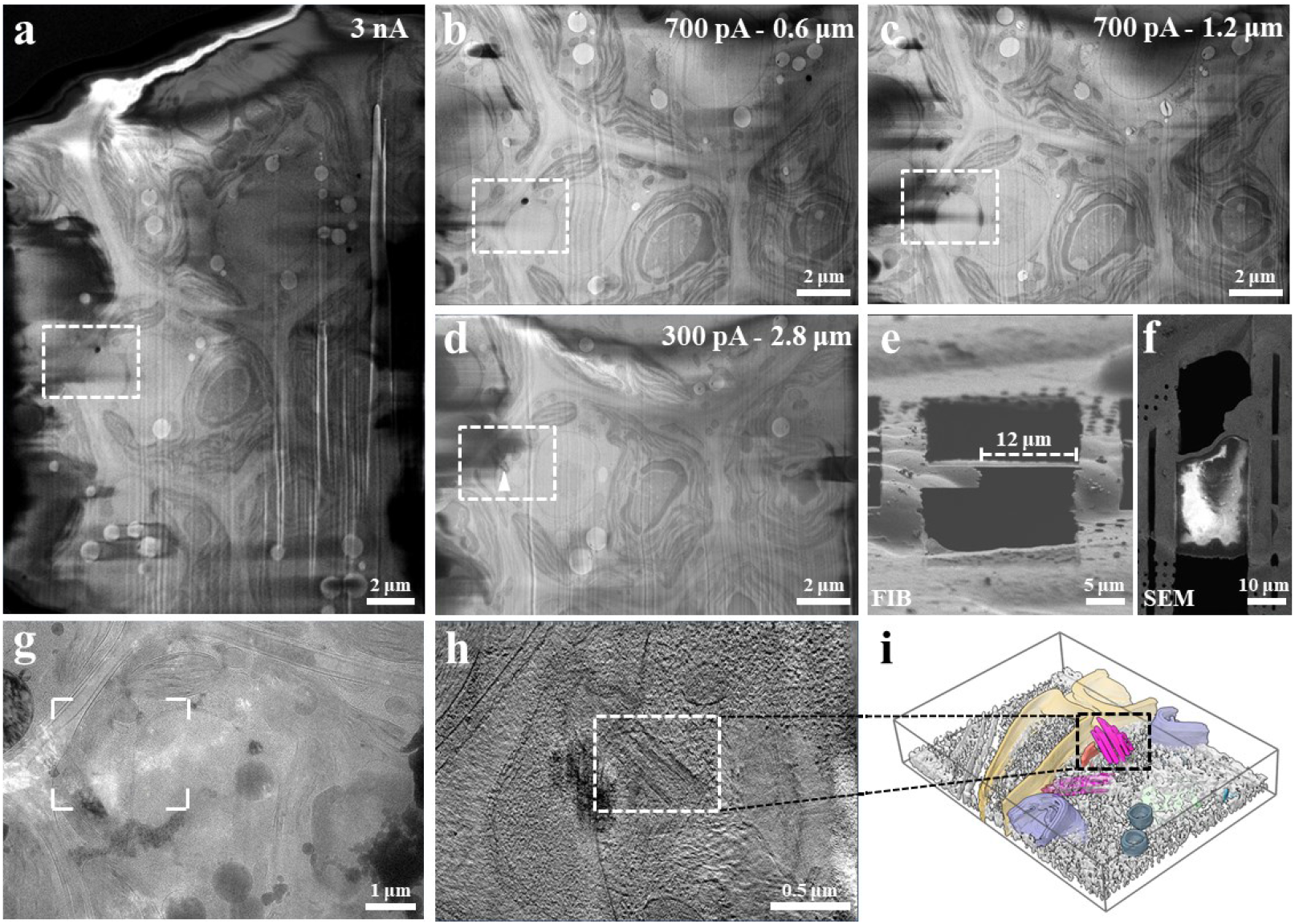
Milling and locating the basal body of *C. reinhardtii* with CSEI. **a**, CSEI of a frozen hydrated *C. reinhardtii* samples after the first milling. A region labeled by a dashed box is recognized as the target basal body. **b**, CESI of the selected region (dashed box) after milling at 0.6 μm depth using a 700 pA current. **c**, CSEI after milling the selected region (dashed box) for a further 0.6 μm (total 1.2 μm) using a 700 pA current. **d**, CSEI after milling the selected region (dashed box) for a further 1.6 μm (total 2.8 μm relative to **a**) using a 300 pA current. The target basal body was pointed at by a white arrow. **e, and f**, The final lamella is shown in FIB and SEM views, respectively. The whole milling procedure is shown in **Supplementary Fig. 18. g**, A low-magnification image of the lamella observed under a 300 kV electron microscope. The target basal body is clearly visible. **h**, A section view of the tomogram with the target basal body. **i**, 3D rendered map of the tomogram shown in **h**. The basal body was pointed by a dashed box in **h** and **i** (purple). Segmented membranes are displayed in different colors. All images of CSEI were collected using Crossbeam 550.

Finally, the prepared lamella (**Fig. 6f**) was examined using cryoET reconstruction. The microtubules in the basal body structure were clearly visible in the tomogram measured with a thickness of ~200 nm (**Fig. 6g-h**). In conclusion, CSEI not only enables the precise localization of specific targets but also provides a serial view of the complete cells gradually. The latter is sometimes important for understanding the relationship between the lamella and the whole cell.

## Discussion

In this work, we introduced CSEI to assist cryoFIB milling, which provided a complete solution for the on-the-fly location without the need for additional hardwares. We discussed the principles of secondary electron imaging on a milled flat surface of frozen hydrated cellular samples. Several key imaging parameters were tested to optimize imaging. Further in-depth studies are required to understand the imaging mechanism of the CSEI. Both our experiments and the reported work on cryoFIB-SEM block face imaging**^8–12^**have demonstrated the feasibility of CSEI. Furthermore, our comparisons also show that CSEI is generally applicable to all tested cryoFIB instruments. While the imaging quality varies among instruments, all tested instruments can meet the basic locating requirements. With further optimization of the CSEI, we believe that most of the tested cryoFIB instruments can achieve better imaging quality.

An important issue in the CSEI is the shadow, which is mostly caused by surface charging. These shadows are either dark (positive surface charge) or bright (negative surface charge), often severely reducing the available imaging area and obstructing the identification of fine structural details. Many factors can cause shadows, including but not limited to the electric conductivity of the bulky sample and the position of the detectors. We also observed that many samples were minimally affected by shadows. The reasons for the generation of shadows or surface charging are still not well understood, and further studies are required.

The cellular contents showed a remarkable contrast in the CSEI. Various membrane structures can be clearly distinguished, and the resolution of the CSEI is sufficient to distinguish densely arranged multilayer membrane structures in the chloroplast stacked thylakoids. The CSEI can also distinguish protein aggregates and contrast variations caused by different protein concentrations. Moreover, the ion beam current, milling flatness, and possible surface damage of the FIB have very little effect on the CSEI, making it a reliable tool for on-the-fly imaging. The current CSEI imaging resolution should meet most of the needs of location requirements during cryoFIB milling.

Overall, CSEI allows us to achieve on-the-fly location during cryoFIB milling without any additional cost. Furthermore, the implementation of CSEI does not have any additional requirements for sample pre-processing and is free of fluorescent labeling. These features significantly enhance cryoFIB to achieve the target of milling arbitrary biological samples. Of course, this technology can also be combined with cryoCLEM technology to meet more diverse location needs.

## Acknowledgements

This work was supported by funds from Tsinghua-Peking Joint Center for Life Sciences, Beijing Frontier Research Center for Biological Structure, and Advanced Innovation Center for Structural Biology. We acknowledge Sihan Wang for providing the mouse liver tissue data, Yaxian Zhao for providing the NRK and HeLa cells, Zhengmao Wang for providing *Chlamydomonas* cells, and Ru Li for providing A431 cells. We acknowledge Xiaomin Li from Tsinghua University and Rui Ma from Thermo Fisher Scientific for technical support. We acknowledge ZEISS Microscopy Customer Center, Beijing lab for providing SEM-FIB facilities. We acknowledge Tsinghua University Branch of China National Center for Protein Sciences Beijing for providing facility supports in computing, cryoEM and SEM-FIB instruments.

## Author contributions

X.L. initialized the project. X.L. and C.L. designed the CSEI experiments. C.L. performed all experiments. L.Z. and Z.Z. assisted C.L. in CSEI experiments and prepared the sample. Y.J. and C.L performed the experiments on Crossbeam 550. X.L. and C.L. wrote the manuscript. All authors revised the manuscript.

## Methods

### *E. coli* cells and cryoEM sample preparation

*E. coli* Rosetta (DE3) cells were grown in LB medium to an OD_600_ of 0.6–0.8 at 37 °C. The cells were collected and resuspended in suspension buffer (50 mM Tris pH 8.0, 300 mM NaCl, and 5% glycerol), and the concentration of the suspension was adjusted to an OD_600_ of 25–40. Subsequently, a drop of 3 μl cell suspension was loaded on a glow-discharged (using a PELCO easiGlow Glow Discharger, Ted Pella Inc) grid (200 mesh gold 1.2/1.3, Quantifoil), and a drop of 2 μl suspension buffer was loaded on the reverse side of the grid. The grid was then blotted from the side with suspension buffer and plunge-frozen using a Leica EM GP (Leica Microsystems). The EM GP was set to a humidity of 75%, a temperature of 25 °C, and a blot time of 6–8 s.

### Mammalian cells and cryoEM sample preparation

HeLa cells were cultured in DMEM (Thermo Fisher Scientific) with 5% CO_2_ at 37 °C. After reaching 70–80% confluence, the cells were digested with 0.25% trypsin-EDTA (Fisher Scientific) for 2 min at 37 °C, washed with PBS, resuspended in DMEM, and diluted to 5 × 10^5^ cells/ml. A431 cells were treated in a similar way, except that the A431 cells were resuspended in PBS rather than in DMEM. The cells were plunge-frozen in the same way as described for *E. coli* above.

### *C. reinhardtii* cells and cryoEM sample preparation

*C. reinhardtii* 21gr cells were cultured as described in the literature**^1^**. Cells were grown to an OD_600_ ~ 2 and harvested by centrifugation for 2 min at 2000 rpm to concentrate the cells 2–8 times. The *C. reinhardtii* cells were plunge-frozen in the same way as described for *E. coli* above.

### *R. sativus* seedling leaf tissue and cryoEM sample preparation

Green leaves of normal growing *R. sativus* seedlings were cut into small pieces and washed 2–3 times in W5 buffer (154 mM NaCl, 25 mM CaCl_2_, 5 mM KCl, 2 mM MES pH 5.7, and 5 mM glucose), then placed in 4% liquid agarose at a temperature of approximately 40 °C. After the agar block solidified completely, the sample was fixed on the sample stage of a vibratome (Leica VT1200 S, Leica Microsystems). The sample tank was filled with W5 buffer before slicing. The sample was sliced with the settings of a slicing frequency of 85 Hz (± 10%), an amplitude of 1 mm, a slicing speed of 0.5 mm/s, and a slicing thickness of 50 μm. The prepared tissue slices were picked using tweezers, transferred to W5 buffer, and frozen on a glow-discharged grid (AG150P 200 mesh Cu, Zhongjingkeyi Technology) using a high-pressure freezer (Leica HPM100, Leica Microsystems). The grid was then transferred to Leica UC7+FC7 (Leica Microsystems) at −150 °C to separate the sapphire and the carrier by volatilizing the protective agent dimethyl pentane. After approximately 20 min, the grids were transferred to a liquid-nitrogen tank for storage.

### Pre-processing before cryoFIB milling

A dual-beam FIB-SEM system (Helios NanoLab DualBeam G3 UC, Thermo Fisher) equipped with a cryo-stage (PP3010T, Quorum) was used to prepare the lamellae.

For the sample vitrified by the plunge-freezing method, the specimen was mounted into an AutoGrid (Thermo Fisher Scientific), loaded into a custom-made sample shuttle, and transferred into the prep stage. A sputter coating (current 5 mA, 60 s) was applied in the prep chamber. The shuttle was then transferred to a cryo-stage. The sample was kept at −180 °C throughout the procedure. Before milling, organometallic platinum deposition was performed on the AutoGrid using the gas injection system (GIS) to reduce radiation damage and curtain effects. The cryo-stage was lowered 4 mm below the eucentric position. The GIS was then turned on for beam-induced Pt deposition of 30 s at 42 °C with an electron beam with an accelerating voltage of 2 kV and a current of 0.4 nA at 100x magnification.

For tissue samples vitrified by high-pressure freezing, the AutoGrid should stay for a longer time of 1–2 h at −150 °C in the prep stage to volatize the protective agent prior to sputter coating (5 mA, 60 s). The first beam-induced Pt deposition was performed following the same procedure as that used for the plunge-frozen specimen. The second Pt deposition was FIB-induced for 20 s at 42 °C, with an accelerating voltage of 30 kV and a current of 33 pA at 100x magnification and a working distance of 4 mm.

### FIB milling and SEM locating

After pretreatment, AutoGrids were transferred to Crossbeam 550 (Carl Zeiss Microscopy GmbH), Helios Nanolab G3 UC (Thermo Fisher Scientific), Aquilos 1 Cryo-FIB (Thermo Fisher Scientific), and Aquilos 2 Cryo-FIB (Thermo Fisher Scientific) to separately execute FIB milling and CSEI. During the entire process, the temperature was maintained at −180 °C.

The entire procedure of FIB milling and SEM locating for *C. reinhardtii* cells was carried out using Crossbeam 550 (**Supplementary Fig. 18**). During cryoFIB milling, an appropriate milling angle was first selected according to the thickness of the sample, which was usually between 13° and 18°. Then, a larger Gallium ion beam current of 3 nA was chosen for the first rough milling with a milling window width of 20 μm under the FIB view. Subsequently, the ion beam current was reduced to 700 pA. Multiple milling steps with a milling depth of 600 nm were performed together with the CSEI. Once the target region was achieved, the sample was milled from the reverse side with a gradually reduced ion beam current, that is, using an ion beam current of 3 nA to 7 μm left, 700 pA to 4 μm left, 300 pA to 2 μm left, and 100 pA to 1 μm left. The milling window was then narrowed to a width of 12 μm using an ion beam current of 50 pA. In the final fine milling step, the two sides of the lamella were polished using an ion beam current of 50 pA.

CSEI in Crossbeam 550 used the settings of an accelerating voltage of 3 kV, an electron beam current of 50 pA, a dwell time of 1.8 μs (scan speed of 5), repetitive scans of 20 times. The final images were merged using images taken by the in-lens and in-chamber detectors with a mixing ratio of between 0.5 and 0.7.

In the test of CSEI by Crossbeam 550, the *E. coli* sample was first tilted to an appropriate angle between 13° and 18° relative to the incident direction of the ion beam. An ion beam current of 700 pA was used for the first rough milling from two sides of the lamella, with a width of 14 μm and a spacing of 3 μm. Subsequently, an ion beam current of 300 pA was used to reduce the thickness to 2 μm. Then the width of the milling window was reduced to 12 μm. The lamella was polished by removing the ~100 nm thickness using an ion beam current of 50 pA ahead of each CSEI test. Before each CSEI test, the focus, astigmatism and other required SEM alignments were performed to ensure imaging quality. Different parameters were tested, including accelerating voltage of 1 kV/ 2 kV/ 3 kV/ 5 kV, electron beam current of 25 pA/ 50 pA/ 100 pA, dwell time of 0.5 μs (scan speed of 3)/ 0.9 μs (scan speed of 4)/ 1.8 μs (scan speed of 5)/ 3.5 μs (scan speed of 6), repetitive scans of 1 time/ 20 times/ 40 times, and working distance of 3.5 mm/ 5 mm/ 7 mm. In-lens and in-chamber detectors were used in all the experiments.

In the test of CSEI by Helios and Aquilos (Aquilos 1 and Aquilos 2), the *E. coli* sample lamella was milled in the same way as described above. The direct alignments should also be adjusted to optimize the imaging conditions ahead of each CSEI test, including the lens alignment, source tilt, and stigmator centering. Focus centering should also be adjusted in Aquilos. Similar imaging conditions of the CSEI were tested as described above for Crossbeam 550, including the accelerating voltage, electron beam current, dwell time, number of repetitive scans (image integration), and working distance. Furthermore, different modes and detectors of Helios and Aquilos were tested, including Mode 1/2 in Helios, Standard/ OptiTilt in Aquilos, ETD/ TLD/ ICE detectors of Helios, ETD/ T2 detectors of Aquilos. All images obtained by CSEI were adjusted for contrast and brightness using ImageJ**^2, 3^**.

### CryoET data collection and reconstruction

CryoET data were collected using a 300 kV Titan Krios electron microscope (Thermo Fisher Scientific) equipped with a GIF quantum energy filter (slit width of 20 eV). Micrographs were recorded with a K3 Summit direct electron detector (Gatan) working in super-resolution mode at a nominal magnification of 19,500, resulting in a calibrated pixel size of 2.261 Å. Tilt series were collected using the bidirectional tilt scheme, first from −13° to −49° and followed from −11° to 39° with an angular increment of 2°, at defocus ranging from −4 to −6 μm by SerialEM**^4^**. A micrograph with eight frames (0.213 s/frame) was recorded at each tilt angle, and the total dose for the tilt series was 90 e/Å^-2^. Beam-induced motion was corrected using MotionCor2**^5^**. The tilt series were aligned and reconstructed using IMOD**^6^**. The final tomograms were processed using IsoNet**^7^**. The segmentation and surface rendering of the density map were performed by Amira (Thermo Fisher Scientific, Mercury Computer Systems), and the 3D rendered figures were prepared using ChimeraX**^8^**.

## Supplementary Figures and legends

**Supplementary Figure 1.**
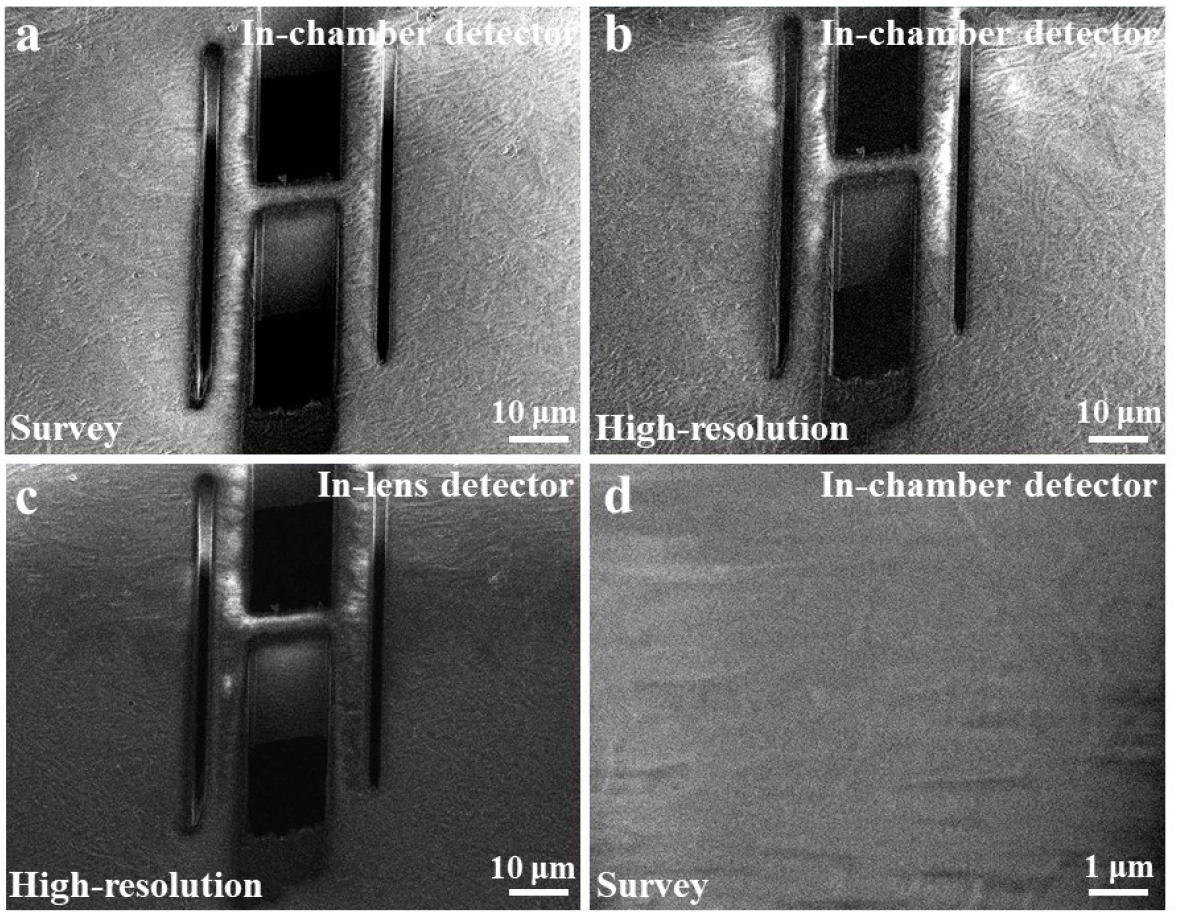
Typical secondary electron images of cryoFIB milled sample under the default settings optimized for topographical imaging using Aquilos 2. **a**, A secondary electron image of a milled *E. coli* sample, imaged using an in-chamber detector with the default settings of the survey mode (Standard mode in Aquilos 2). **b**, A secondary electron image of the same sample as that in **a**, imaged using an in-chamber detector with the default settings of the high-resolution mode (OptiTilt mode in Aqulios2). **c**, A secondary electron image of the same sample as that in **a**, imaged using an in-lens detector (T2 in Aquilos 2) with the default settings of the high-resolution mode (OptiTilt in Aquilos 2). **d**, The secondary electron image of a cryoFIB milled surface of a frozen hydrated *E. coli* sample, imaged using the same imaging conditions as those in **a**. The image shows the nearly invisible contrast of the cells.

**Supplementary Figure 2.**
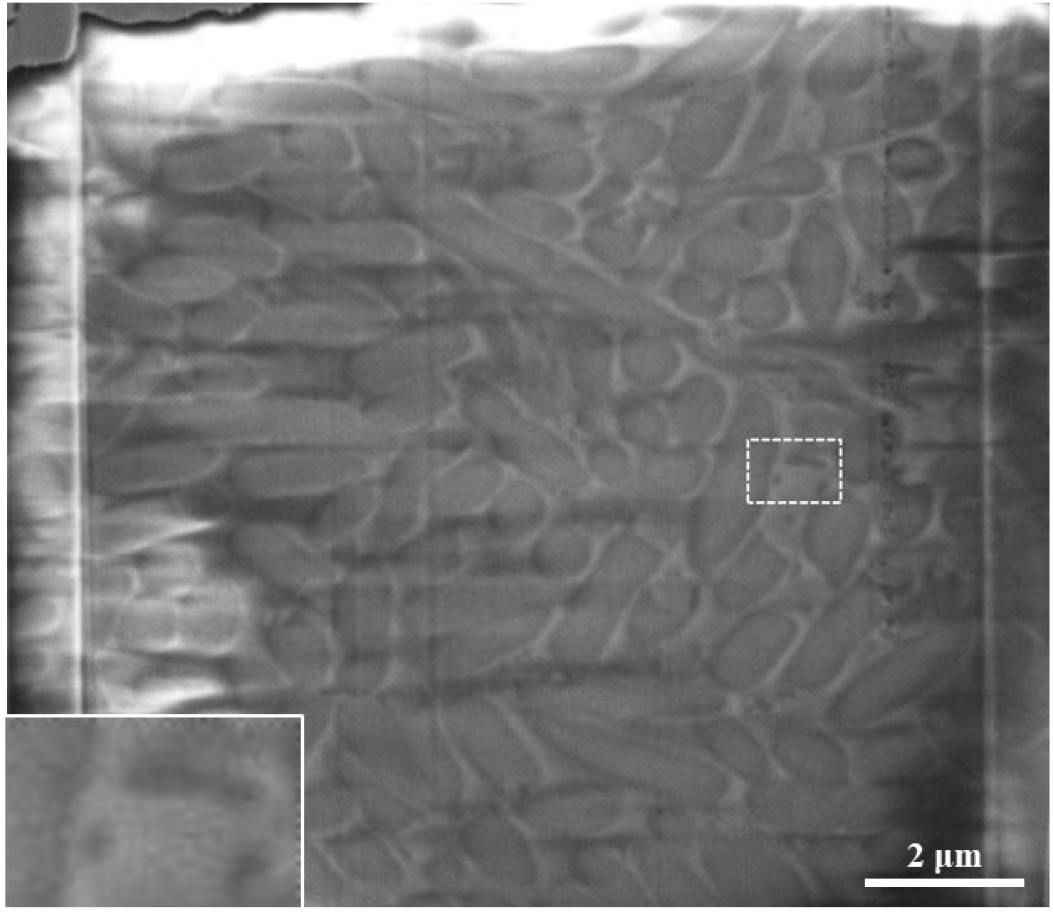
*E. coli* cells visualized using an in-chamber detector named ICE in Helios. The ICE detector in Helios belongs to the in-chamber detector. The imaging quality of the ICE detector for CSEI was not as good as the other detectors in our test. Therefore, we did not include this detector in the comparison of Fig. 2 and just showed a typical image acquired by the ICE detector here. A small rectangle region is magnified and inset at the bottom left. The image was recorded at 2 kV in the survey mode (“mode 1” in Helios) using the same imaging parameters as those of **Fig. 2a** and **e**.

**Supplementary Figure 3.**
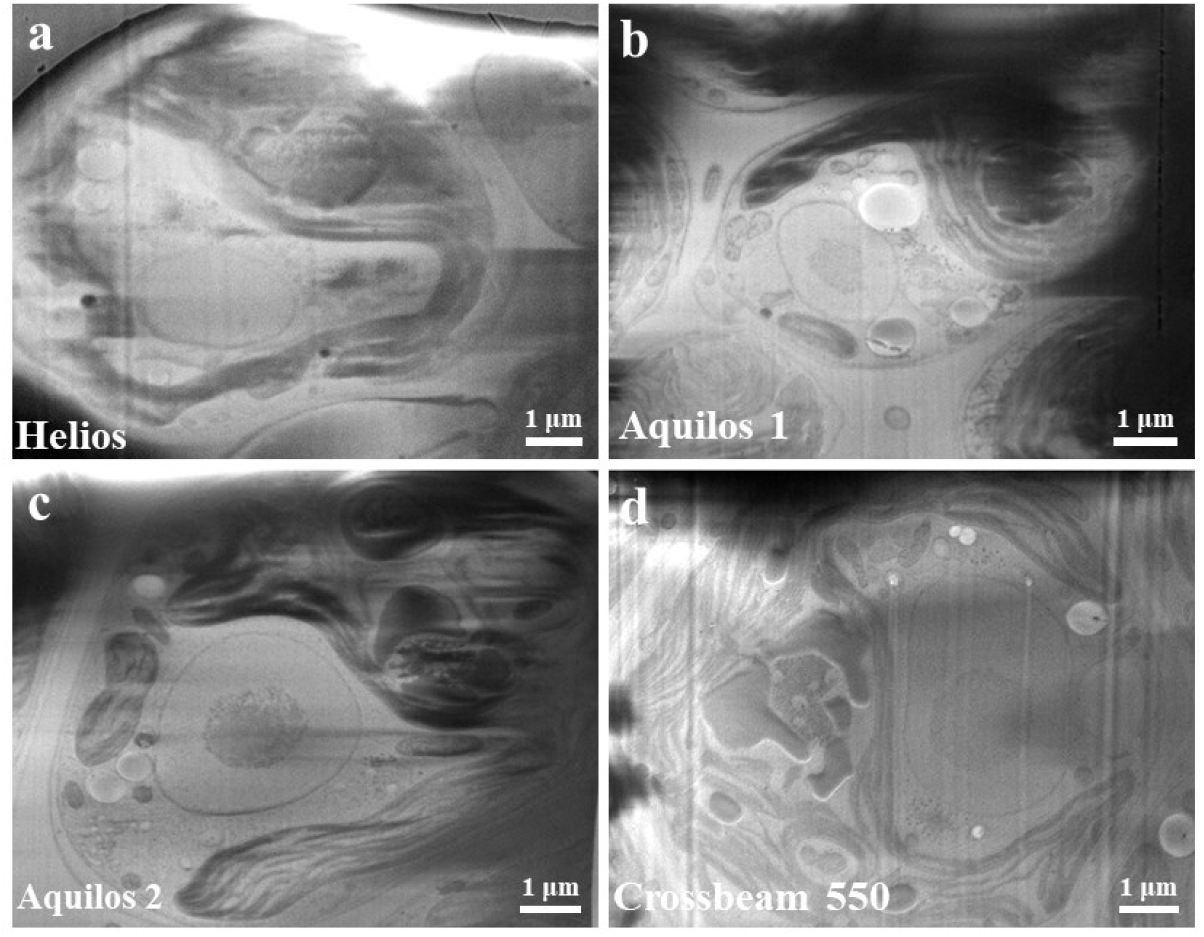
*C. reinhardtii* cells visualized using different cryoFIB instruments. **a**, An image acquired on Helios. **b**, An image acquired on Aquilos 1. **c**, An image acquired on Aquilos 2. The imaging for **a**, **b,** and **c** used the same settings with a voltage of 2 kV, electron beam current of 50 pA, a dwell time of 1 μs and repetitive scans of 20 times. **d**, An image acquired on Crossbeam 550 with a voltage of 3 kV, an electron beam current of 50 pA, a dwell time of 1.8 μs (scan speed of 5), and repetitive scans of 20 times. All the images were acquired on the cryoFIB milled surface of frozen hydrated *C. reinhardtii* cells. Crossbeam 550 is from a different vendor than the other three instruments, so some settings are slightly different to achieve the best image quality.

**Supplementary Figure 4.**
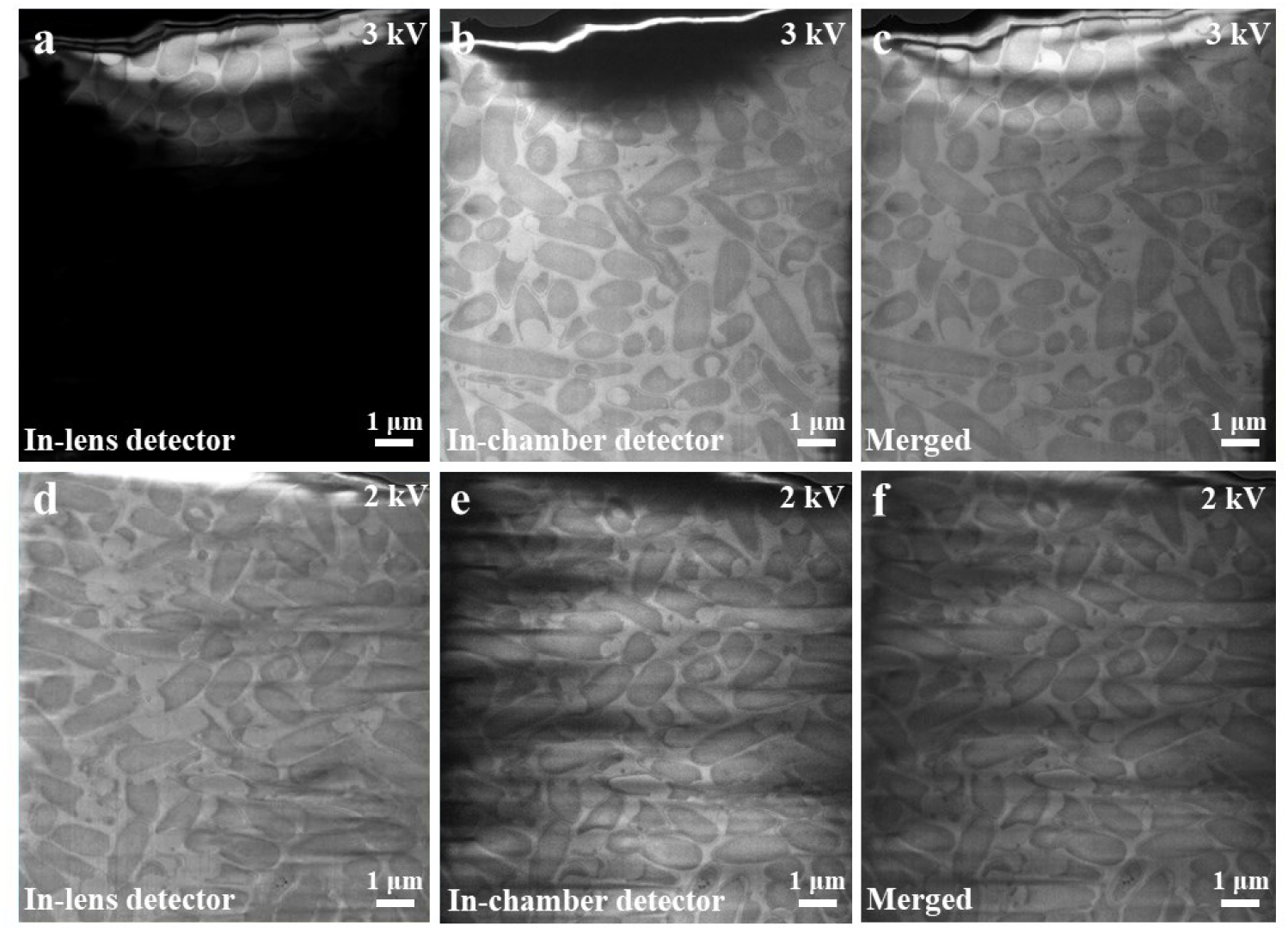
*E. coli* cells visualized by different detectors of Crossbeam 550. **a,** and **b,** Images acquired by the in-lens and in-chamber detectors, respectively, at a voltage of 3 kV. **c**, A merged image of **a** and **b**. The image **a** was significantly influenced by the shadow, which was frequently observed when using the in-lens detector on the Crossbeam 550. The merged image shows a complete view of the imaged area, demonstrating that the shadows appearing in **a** and **b** are complementary. **d,** and **e,** Images acquired by the in-lens and in-chamber detectors, respectively, at a voltage of 2 kV. **f**, A merged image of **d** and **e**. Under the lower voltage of 2 kV, the shadow issue that appeared on the in-lens detector disappeared. The reason is still not clear. All the images were acquired on the cryoFIB milled surface of frozen hydrated *E. coli* cells.

**Supplementary Figure 5.**
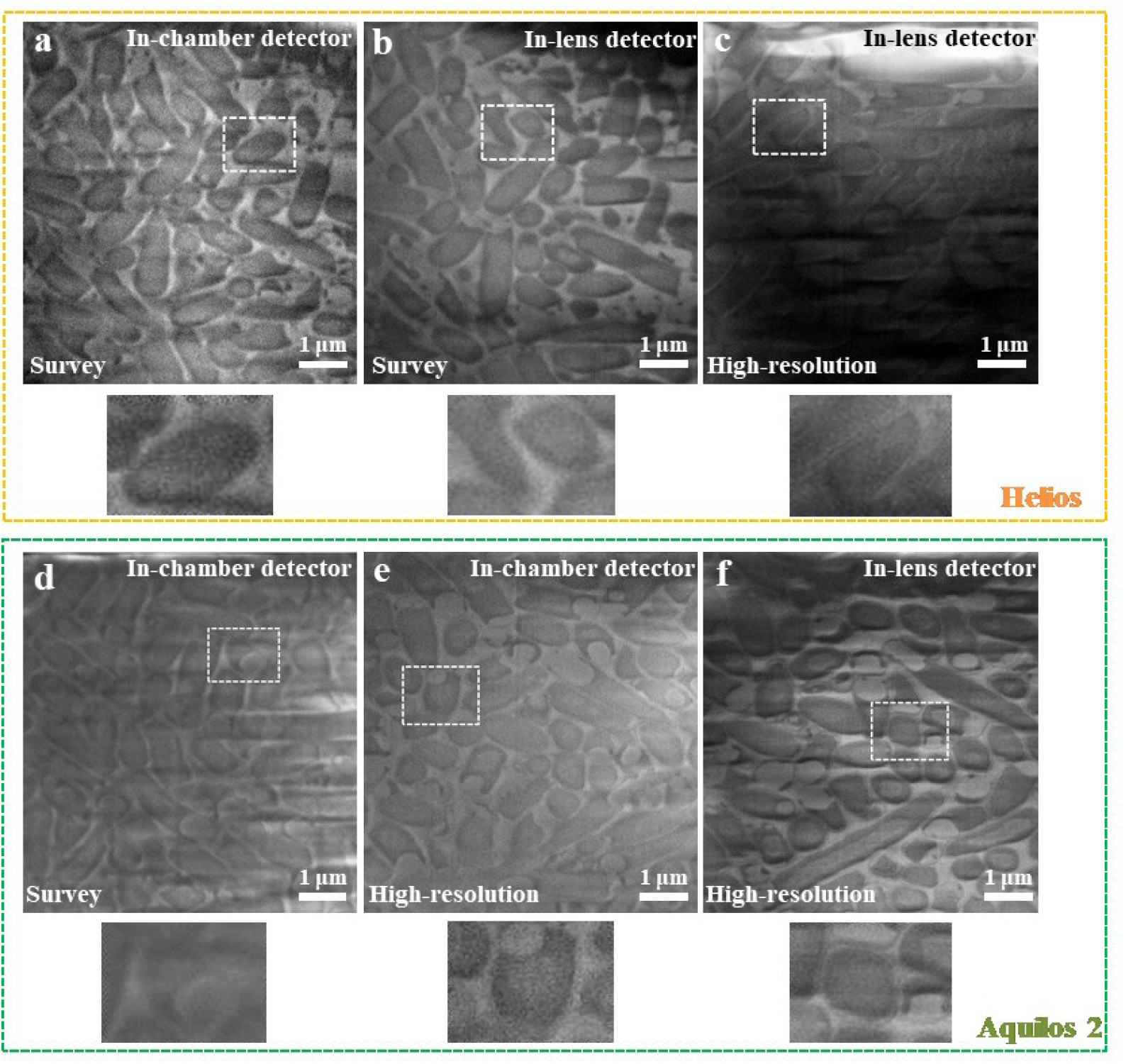
Comparison of the survey and high-resolution mode in Helios and Aquilos 2 for the purpose of CSEI. **a**-**c**, Images acquired on Helios using different detectors and modes as labeled in the figure. **d**-**f**, Images acquired on Aquilos 2 using different detectors and modes as labeled in the figure. All images were recorded using a voltage of 2 kV, an electron beam current of 50 pA, a dwell time of 1 μs, and repetitive scans of 20 times. A small rectangle region in each figure is magnified and shown below the corresponding image. All the images were acquired on the cryoFIB milled surface of frozen hydrated *E. coli* cells.

**Supplementary Figure 6.**
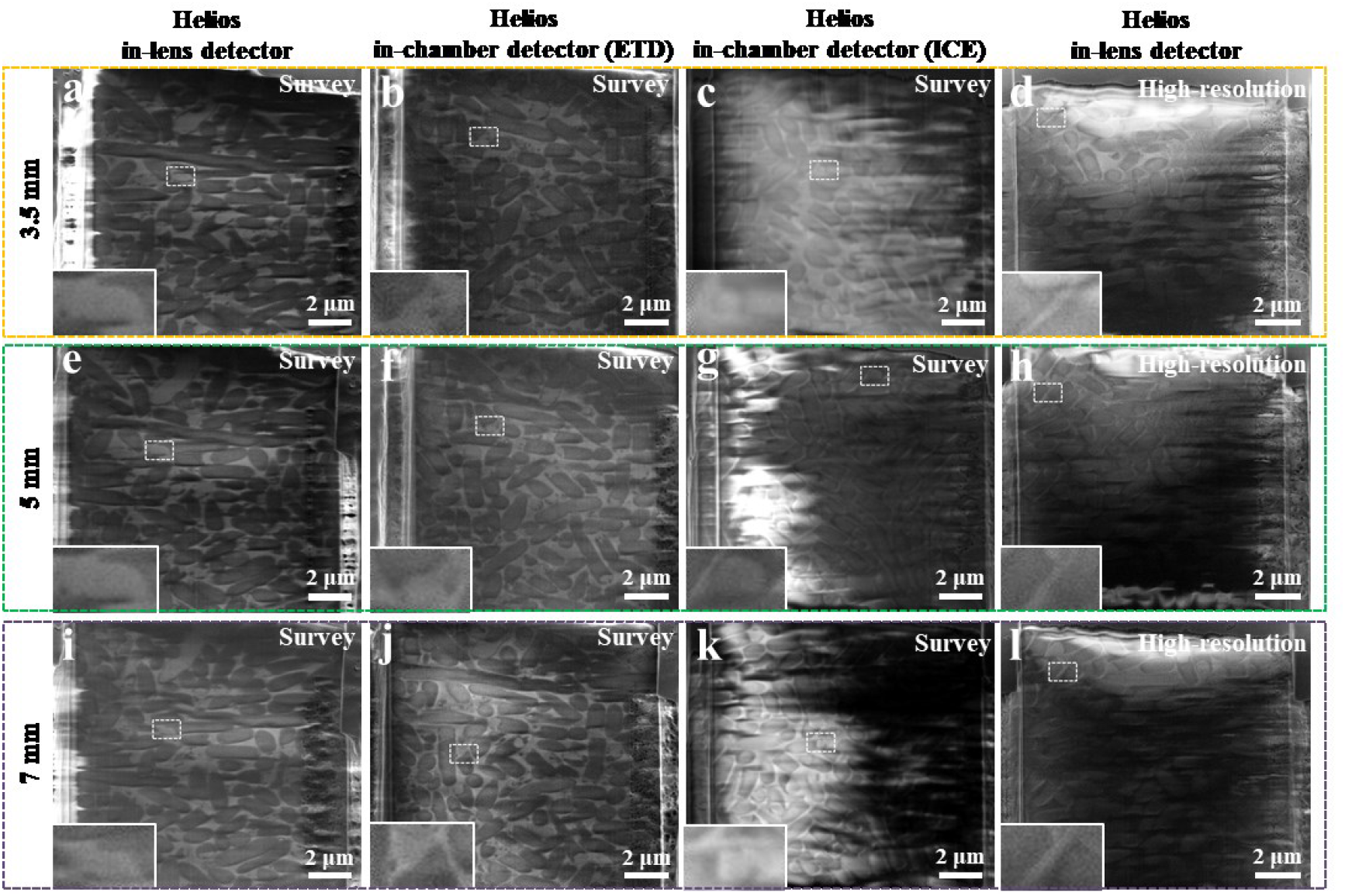
Comparison of the influence of the working distance on CSEI for Helios. **a-d**, Typical images acquired at a working distance of 3.5 mm using different detectors and imaging mode settings as labeled in each figure. **e-h**, Typical images acquired at a working distance of 5 mm using different detectors and imaging mode settings as labeled in each figure. **i-l**, Typical images acquired at a working distance of 7 mm using different detectors and imaging mode settings as labeled in each figure. All images were recorded using a voltage of 2 kV, an electron beam current of 50 pA, a dwell time of 1 μs, and repetitive scans of 20 times. A small rectangle region in each figure is magnified and inset at the bottom left of the corresponding image. All the images were acquired on the cryoFIB milled surface of frozen hydrated *E. coli* cells.

**Supplementary Figure 7.**
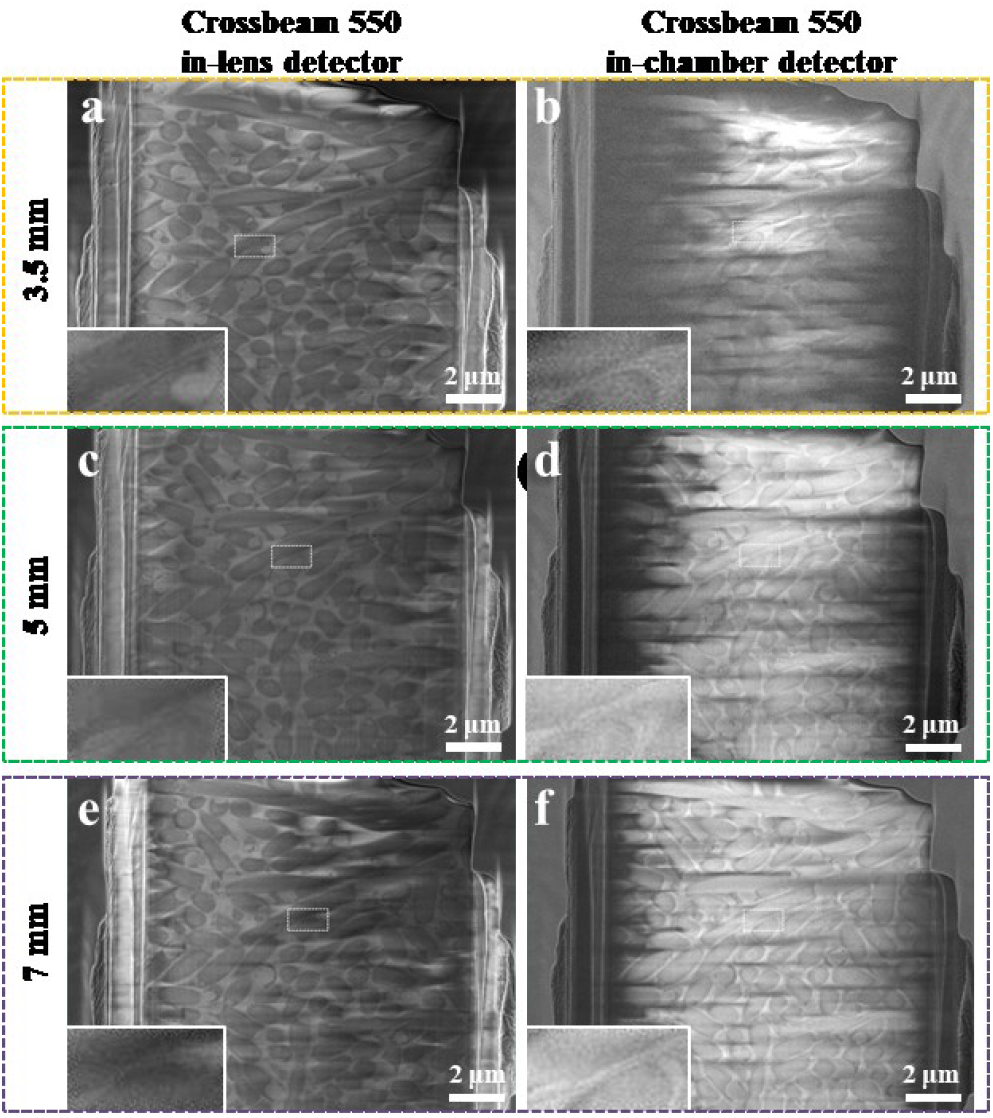
Comparison of the influence of the working distance on CSEI for Crossbeam 550. **a-b**, Typical images acquired at a working distance of 3.5 mm using different detectors and imaging mode settings as labeled in each figure. **c-d**, Typical images acquired at a working distance of 5 mm using different detectors and imaging mode settings as labeled in each figure. **e-f**, Typical images acquired at a working distance of 7 mm using different detectors and imaging mode settings as labeled in each figure. All images were recorded using a voltage of 2 kV, an electron beam current of 50 pA, a dwell time of 1.8 μs (scan speed of 5), and repetitive scans of 20 times. A small rectangle region in each figure is magnified and inset at the bottom left of the corresponding image. All the images were acquired on the cryoFIB milled surface of frozen hydrated *E. coli* cells.

**Supplementary Figure 8.**
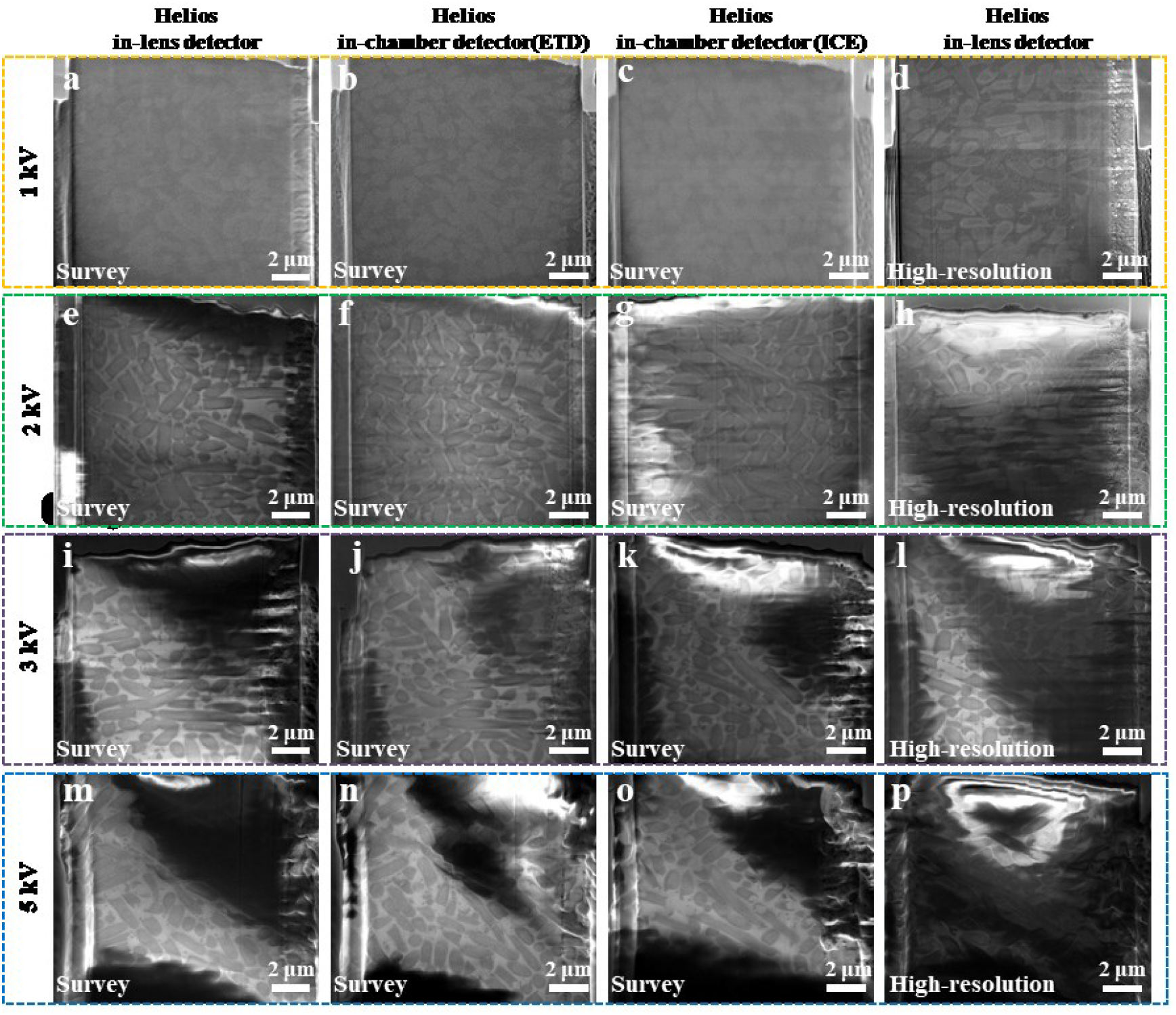
Comparison of the influence of the voltage on CSEI for Helios. **a-d**, Typical images acquired at a voltage of 1 kV using different detectors and imaging mode settings as labeled in each figure. **e-h**, Typical images acquired at a voltage of 2 kV using different detectors and imaging mode settings as labeled in each figure. **i-l**, Typical images acquired at a voltage of 3 kV using different detectors and imaging mode settings as labeled in each figure. **m-p**, Typical images acquired at a voltage of 5 kV using different detectors and imaging mode settings as labeled in each figure. All images were recorded using an electron beam current of 50 pA, a dwell time of 1 μs, and repetitive scans of 20 times. All the images were acquired on the cryoFIB milled surface of frozen hydrated *E. coli* cells.

**Supplementary Figure 9.**
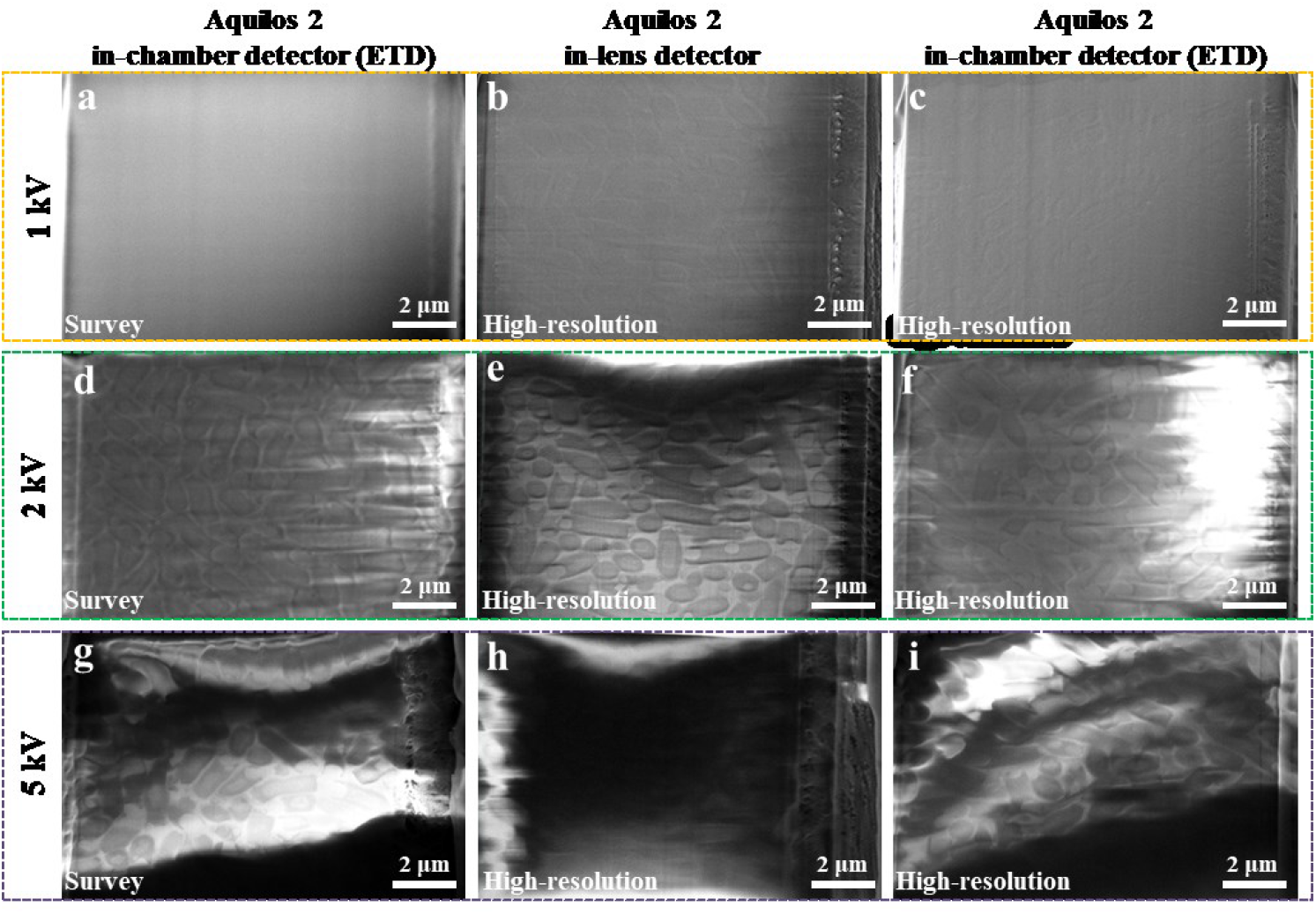
Comparison of the influence of the voltage on CSEI for Aquilos 2. **a-c**, Typical images acquired at a voltage of 1 kV using different detectors and imaging mode settings as labeled in each figure. **d-f**, Typical images acquired at a voltage of 2 kV using different detectors and imaging mode settings as labeled in each figure. **g-i**, Typical images acquired at a voltage of 5 kV using different detectors and imaging mode settings as labeled in each figure. All images were recorded using an electron beam current of 50 pA, a dwell time of 1 μs, and repetitive scans of 20 times. All the images were acquired on the cryoFIB milled surface of frozen hydrated *E. coli* cells.

**Supplementary Figure 10.**
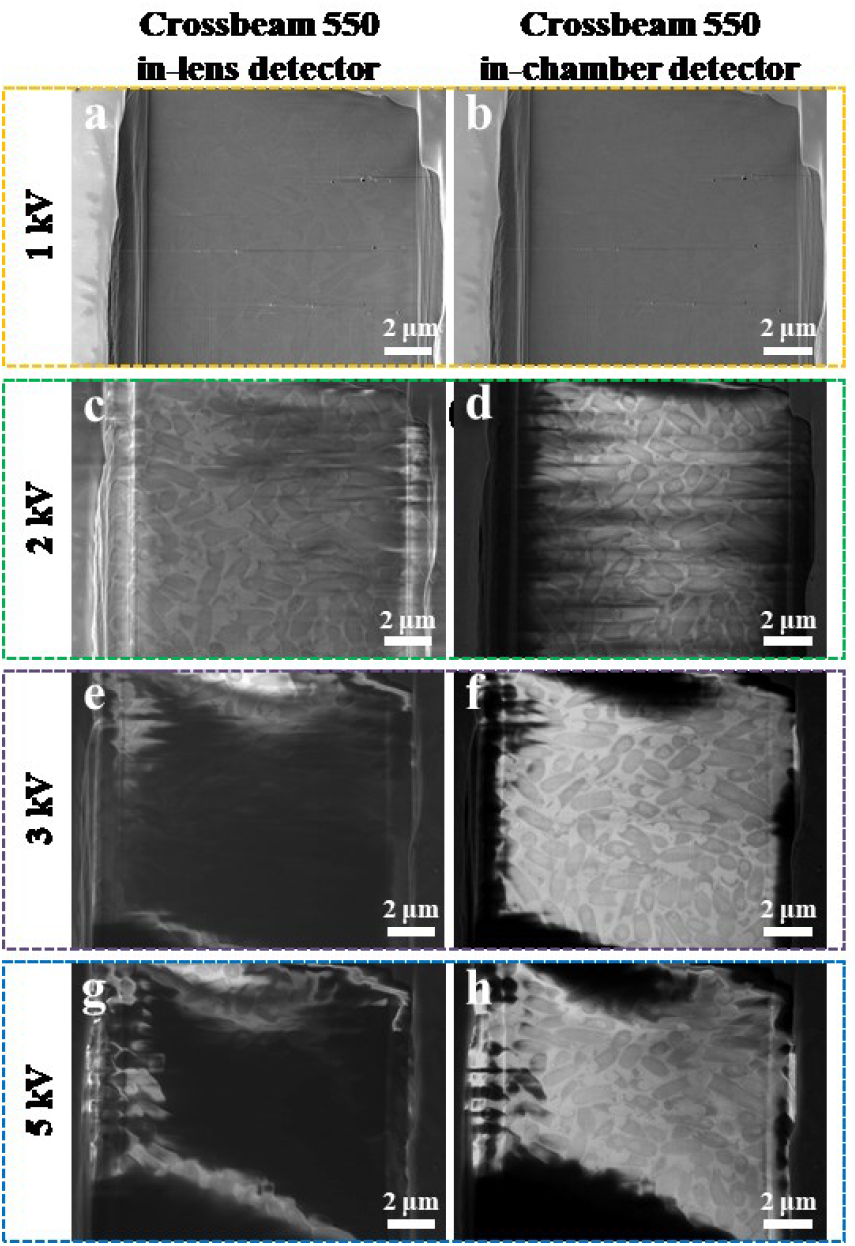
Comparison of the influence of the voltage on CSEI for Crossbeam 550. **a-b**, Typical images acquired at a voltage of 1 kV using different detectors as labeled on the top. **c-d**, Typical images acquired at a voltage of 2 kV using different detectors as labeled on the top. **e-f**, Typical images acquired at a voltage of 3 kV using different detectors as labeled on the top. **g-h**, Typical images acquired at a voltage of 5 kV using different detectors as labeled on the top. Crossbeam 550 only uses a single imaging mode and does not distinguish between the survey and high-resolution mode as other instruments tested in the present work. All images were recorded using an electron beam current of 50 pA, a dwell time of 1.8 μs (scan speed of 5), and repetitive scans of 20 times. All the images were acquired on the cryoFIB milled surface of frozen hydrated *E. coli* cells.

**Supplementary Figure 11.**
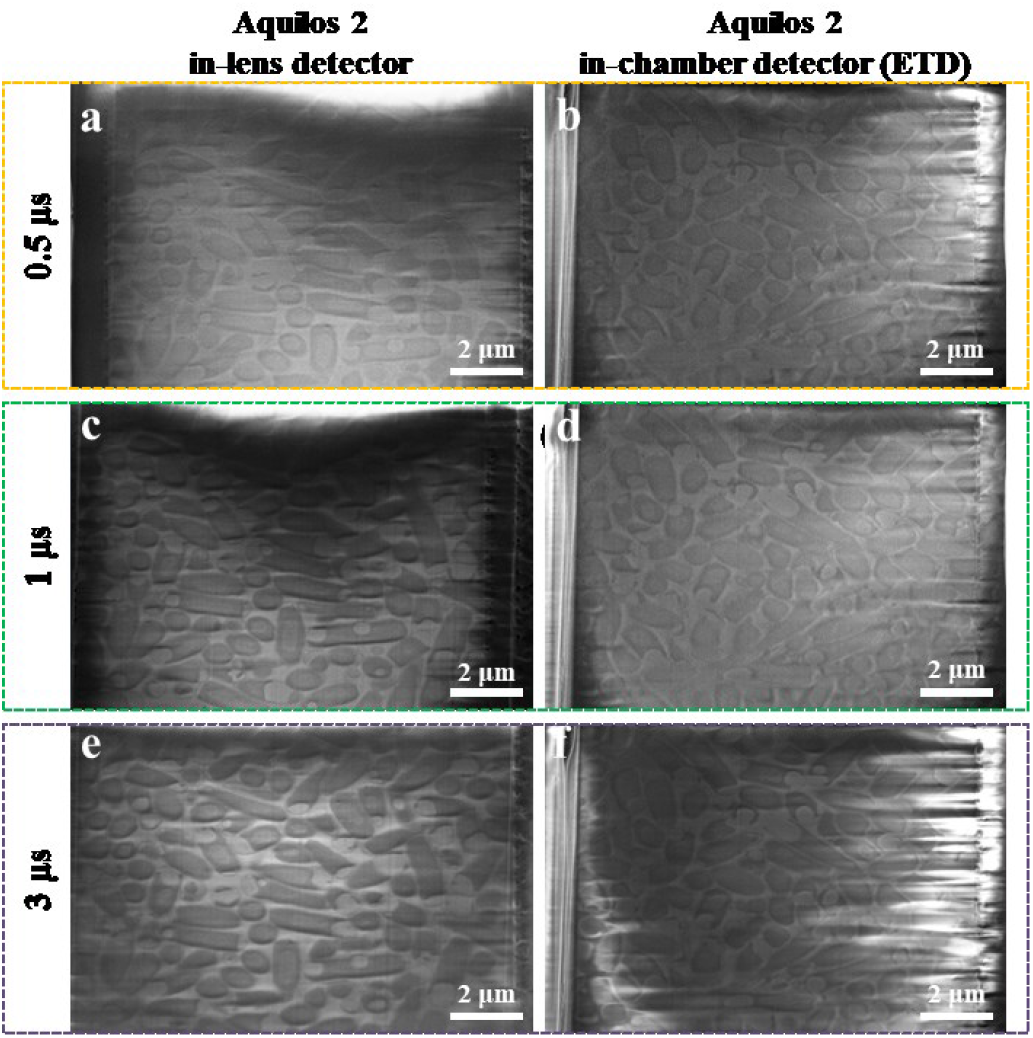
Comparison of the influence of the dwell time on CSEI for Aquilos 2. **a-b**, Typical images acquired at a dwell time of 0.5 μs using different detectors as labeled on the top. **c-d**, Typical images acquired at a dwell time of 1 μs using different detectors as labeled on the top. **e-f**, Typical images acquired at a dwell time of 3 μs using different detectors as labeled on the top. All images were recorded using a voltage of 2 kV, an electron beam current of 50 pA, and repetitive scans of 20 times in the high-resolution mode (OptiTilt). All the images were acquired on the cryoFIB milled surface of frozen hydrated *E. coli* cells.

**Supplementary Figure 12.**
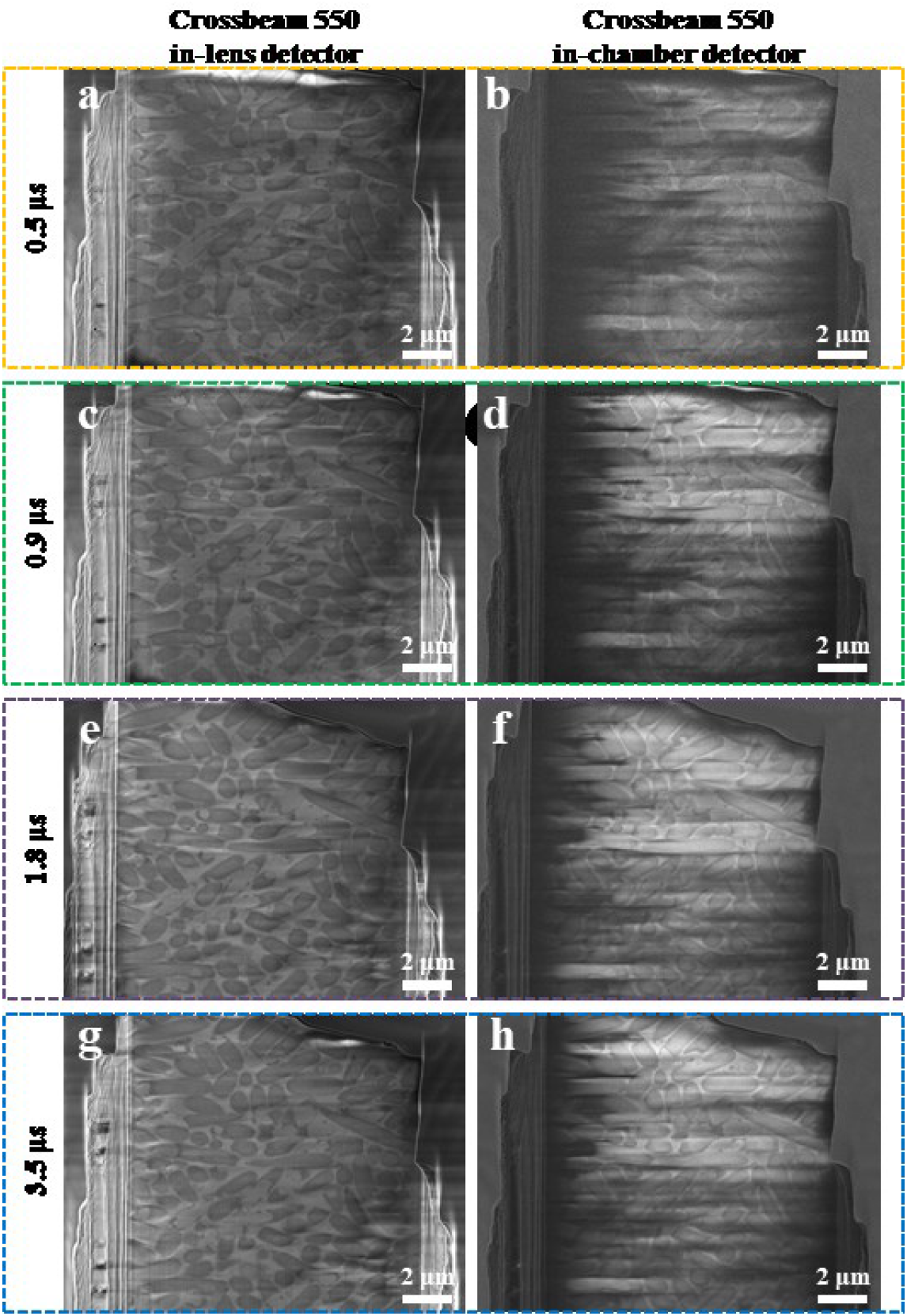
Comparison of the influence of the dwell time on CSEI for Crossbeam 550. **a-b**, Typical images acquired at a dwell time of 0.5 μs (scan speed of 3) using different detectors as labeled on the top. **c-d**, Typical images acquired at a dwell time of 0.9 μs (scan speed of 4) using different detectors as labeled on the top. **e-f**, Typical images acquired at a dwell time of 1.8 μs (scan speed of 5) using different detectors as labeled on the top. **g-h**, Typical images acquired at a dwell time of 3.5 μs (scan speed of 6) using different detectors as labeled on the top. All images were recorded using a voltage of 2 kV, an electron beam current of 50 pA, and repetitive scans of 20 times. All the images were acquired on the cryoFIB milled surface of frozen hydrated *E. coli* cells.

**Supplementary Figure 13.**
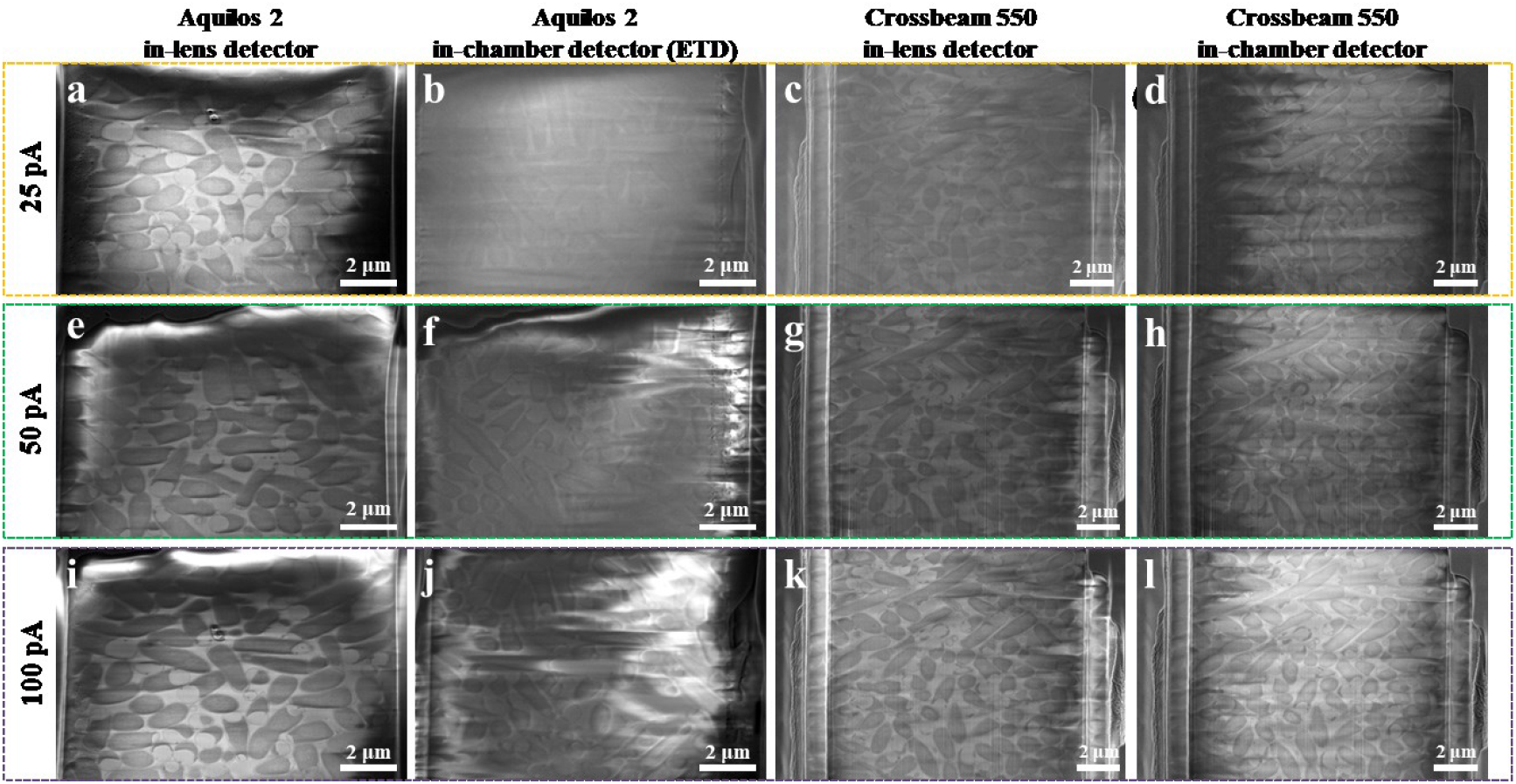
Comparison of the influence of the electron beam current on CSEI. **a-d**, Typical images acquired at an electron beam current of 25 pA using different detectors and instruments as labeled on the top. **e-h**, Typical images acquired at an electron beam current of 50 pA using different detectors and instruments as labeled on the top. **i-l**, Typical images acquired at an electron beam current of 100 pA using different detectors and instruments as labeled on the top. Images were recorded on Aquilos 2 using a voltage of 2 kV, a dwell time of 1 μs, and repetitive scans of 20 times in the high-resolution mode (OptiTilt). Images were recorded on Crossbeam 550 using a voltage of 2 kV, a dwell time of 1.8 μs (scan speed of 5), and repetitive scans of 20 times. All the images were acquired on the cryoFIB milled surface of frozen hydrated *E. coli* cells.

**Supplementary Figure 14.**
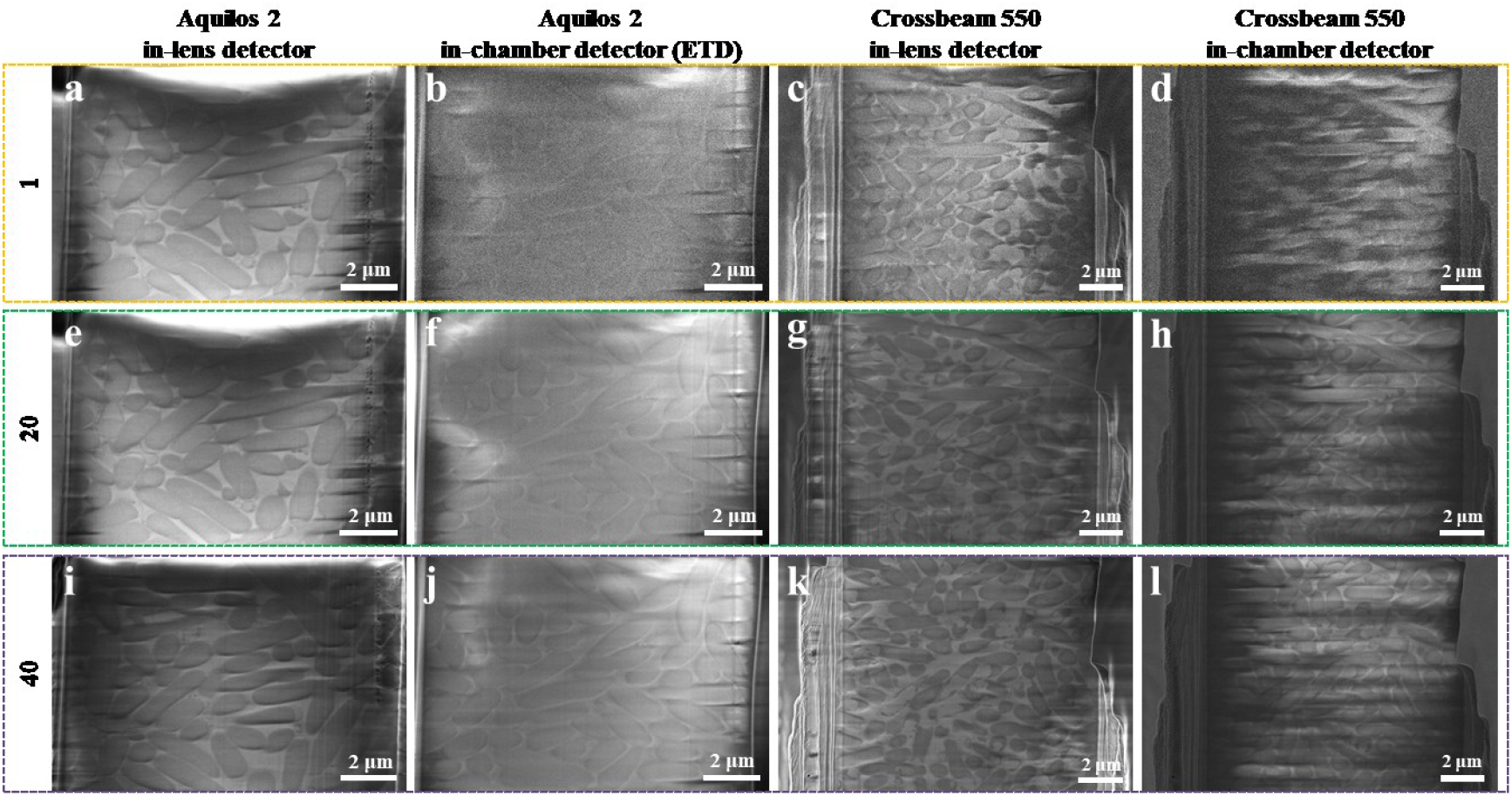
Comparison of the influence of the number of repetitive scans on CSEI. **a-d**, Typical images acquired with repetitive scans of 1 time using different detectors and instruments as labeled on the top. **e-h**, Typical images acquired with repetitive scans of 20 times using different detectors and instruments as labeled on the top. **i-l**, Typical images acquired with repetitive scans of 40 times using different detectors and instruments as labeled on the top. Images were recorded on Aquilos 2 using a voltage of 2 kV, a dwell time of 1 μs, and an electron beam current of 50 pA in the high-resolution mode (OptiTilt). Images were recorded on Crossbeam 550 using a voltage of 2 kV, a dwell time of 1.8 μs (scan speed of 5), and an electron beam current of 50 pA. All the images were acquired on the cryoFIB milled surface of frozen hydrated *E. coli* cells.

**Supplementary Figure 15.**
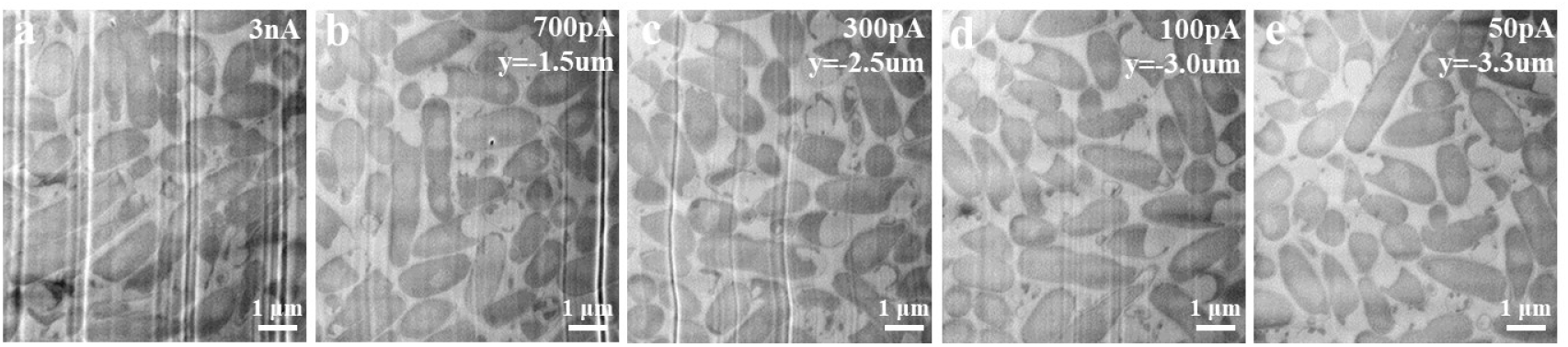
CSEI on the surface milling by different ion beam currents with severe curtaining issue. Sample surfaces of frozen hydrated *E. coli,* milled by different ion beam currents as shown on the top right and visualized by CSEI. Images were recorded on Crossbeam 550 using a voltage of 3 kV, a dwell time of 1.8 μs (scan speed of 5), an electron beam current of 50 pA, and repetitive scans of 20 times.

**Supplementary Figure 16.**
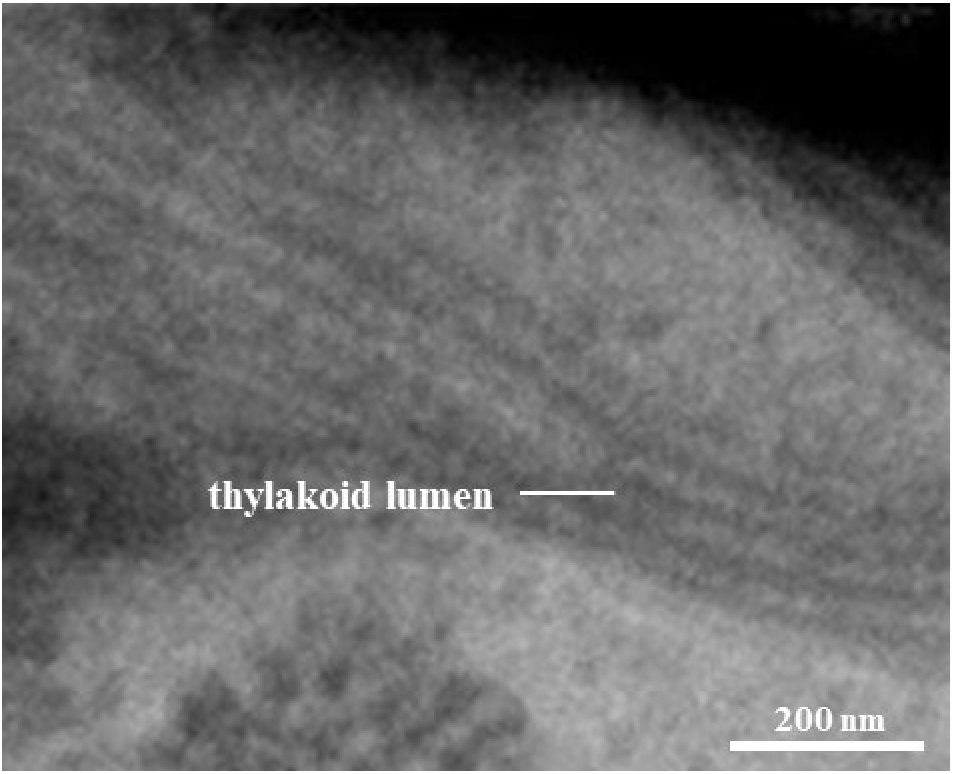
CSEI visualization of stacked membranes in the thylakoid lumen. The densely stacked thylakoids of grana in chloroplasts can be distinguished clearly. The *C. reinhardtii* cell was cryoFIB milled and observed using Crossbeam 550. The image was recorded using a voltage of 3 kV, a dwell time of 1.8 μs (scan speed of 5), an electron beam current of 50 pA, and repetitive scans of 20 times.

**Supplementary Figure 17.**
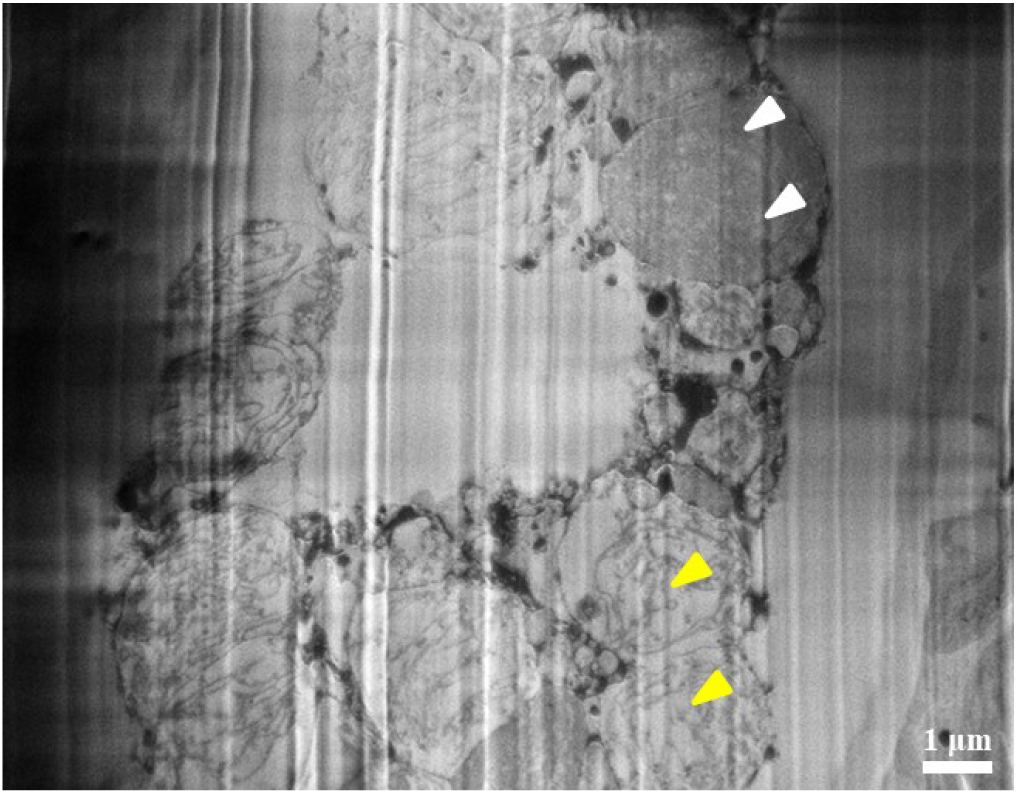
A *R. sativus* tissue sample visualized by CSEI. White arrows point to intact *R. sativus* cells containing chloroplast matrix proteins. Yellow arrows point to *R. sativus* cells that may lose chloroplast matrix proteins, which are lighter in gray level. The loss of chloroplast matrix proteins might be due to the mechanical damage during the sample preparation by a vibratory microtome.

**Supplementary Figure 18.**
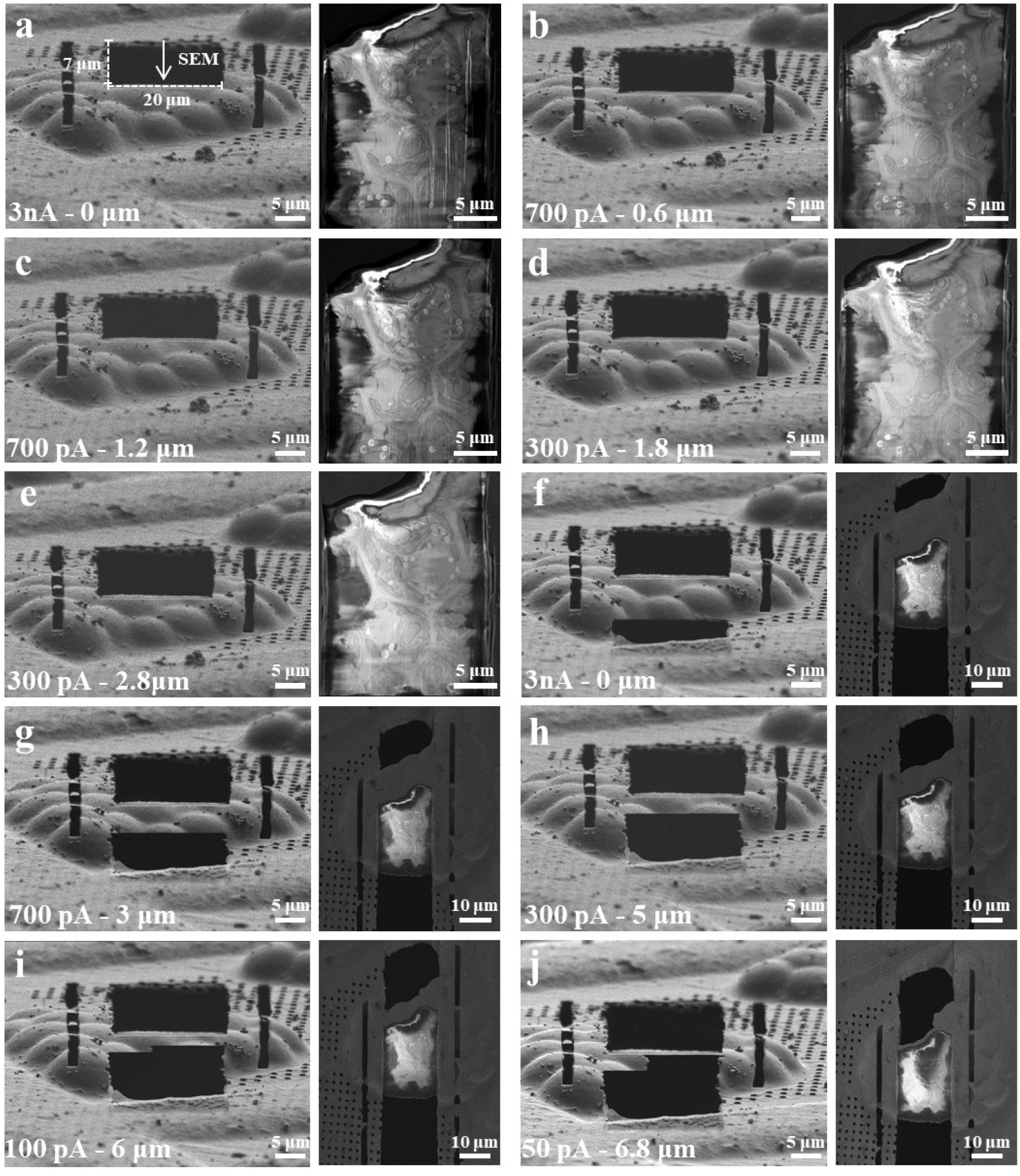
Illustration of a complete milling workflow with on-the-fly locating of the basal body in lamella. The frozen hydrated *C. reinhardtii* cells were used for the test. In each panel, a pair of images under FIB (left) and SEM (right) view is shown. The ion beam current and accumulated milling depth is shown on the bottom of each FIB image. **a**, Initial milling in a window of 20 μm width and 7 μm height under FIB view, and CSEI was performed on the milled surface (white arrow). **b**, **c**, **d,** and **e**, Multiple milling on one side of the sample to locate the target basal body. **f**, The first coarse milling on another side relative to **a**. **g** and **h**, Multiple milling to further reduce the thickness. **i** and **j**, Multiple milling with reduced width to focus on the region with the target basal body. Images were recorded on Crossbeam 550, and CSEI was performed with a voltage of 3 kV, a dwell time of 1.8 μs (scan speed of 5), an electron beam current of 50 pA, and repetitive scans of 20 times.

## Supplementary Tables and Legends

**Supplementary Table 1.** Trade names of detectors on different cryoFIB instruments.

|  | Helios | Aquilos 1 | Aquilos 2 | Crossbeam 550 |
| --- | --- | --- | --- | --- |
| In-lens detector | TLD<sup>a</sup> | T2 | T2 | Inlens |
| In-chamber detector (ETD) | ETD<sup>b</sup> | ETD<sup>b</sup> | ETD<sup>b</sup> | SE2<sup>c</sup> |
| In-chamber detector (ICE) | ICE<sup>d</sup> |  |  |  |

<sup>a</sup>The Through Lens Detector (TLD).
<sup>b</sup>The Everhart Thornley Detector (ETD) is permanently mounted in the chamber over and to one side of the sample.
<sup>c</sup>The SE2 detector is an Everhart Thornley type detector.
<sup>d</sup>The In Chamber Electronics (ICE) detector is mounted near the end of the ion column. Only Helios has the ICE detector.

**Supplementary Table 2.** Trade names of modes on different cryoFIB instruments. Crossbeam 550 does not differentiate between the survey mode and the high-resolution mode.

|  | Helios | Aquilos 1 | Aquilos 2 |
| --- | --- | --- | --- |
| Survey mode | Mode 1<sup>a</sup> | Standard | Standard |
| High-resolution mode | Mode 2<sup>b</sup> | OptiTilt | OptiTilt |

<sup>a</sup>Mode 1 is the default survey mode in Helios and is essential for navigation and assessment of the sample during the FIB milling process. In Mode 1, the immersion lens is switched off.
<sup>b</sup>In Mode 2, the immersion lens is switched on.

## References

1. Lucić, V. et al. Multiscale imaging of neurons grown in culture: from light microscopy to cryo-electron tomography. J Struct Biol 160, 146–156 (2007).

2. van Driel, L.F., Valentijn, J.A., Valentijn, K.M., Koning, R.I. & Koster, A.J. Tools for correlative cryo-fluorescence microscopy and cryo-electron tomography applied to whole mitochondria in human endothelial cells. Eur J Cell Biol 88, 669–684 (2009).

3. Chang, Y.W. et al. Correlated cryogenic photoactivated localization microscopy and cryo-electron tomography. Nat Methods 11, 737–739 (2014).

4. Tuijtel, M.W., Koster, A.J., Jakobs, S., Faas, F.G.A. & Sharp, T.H. Correlative cryo super-resolution light and electron microscopy on mammalian cells using fluorescent proteins. Sci Rep 9, 1369 (2019).

5. Liu, B. et al. Three-dimensional super-resolution protein localization correlated with vitrified cellular context. Sci Rep 5, 13017 (2015).

6. Dahlberg, P.D. et al. Cryogenic single-molecule fluorescence annotations for electron tomography reveal in situ organization of key proteins in Caulobacter. Proc Natl Acad Sci U S A 117, 13937–13944 (2020).

7. Dahlberg, P.D. & Moerner, W.E. Cryogenic Super-Resolution Fluorescence and Electron Microscopy Correlated at the Nanoscale. Annu Rev Phys Chem 72, 253–278 (2021).

8. Schertel, A. et al. Cryo FIB-SEM: volume imaging of cellular ultrastructure in native frozen specimens. J Struct Biol 184, 355–360 (2013).

9. Sviben, S. et al. A vacuole-like compartment concentrates a disordered calcium phase in a key coccolithophorid alga. Nat Commun 7, 11228 (2016).

10. Vidavsky, N. et al. Cryo-FIB-SEM serial milling and block face imaging: Large volume structural analysis of biological tissues preserved close to their native state. J Struct Biol 196, 487–495 (2016).

11. Spehner, D. et al. Cryo-FIB-SEM as a promising tool for localizing proteins in 3D. J Struct Biol 211, 107528 (2020).

12. Zhu, Y. et al. Serial cryoFIB/SEM Reveals Cytoarchitectural Disruptions in Leigh Syndrome Patient Cells. Structure 29, 82–87.e83 (2021).

13. Stokes, D. Principles and practice of variable pressure/environmental scanning electron microscopy (VP-ESEM). (John Wiley & Sons, 2008).

14. Joy, D.C. & Joy, C.S. Study of the Dependence of E2 Energies on Sample Chemistry. Microsc Microanal 4, 475–480 (1998).

15. Hayles, M.F., Stokes, D.J., Phifer, D. & Findlay, K.C. A technique for improved focused ion beam milling of cryo-prepared life science specimens. J Microsc 226, 263–269 (2007).

16. Mackinder, L.C. et al. A repeat protein links Rubisco to form the eukaryotic carbon-concentrating organelle. Proc Natl Acad Sci U S A 113, 5958–5963 (2016).

17. Freeman Rosenzweig, E.S. et al. The Eukaryotic CO(2)-Concentrating Organelle Is Liquid-like and Exhibits Dynamic Reorganization. Cell 171, 148–162.e119 (2017).

18. Erdmann, P.S. et al. In situ cryo-electron tomography reveals gradient organization of ribosome biogenesis in intact nucleoli. Nat Commun 12, 5364 (2021).

19. Wingfield, J.L. & Lechtreck, K.F. Chlamydomonas Basal Bodies as Flagella Organizing Centers. Cells 7 (2018).

20. Geimer, S. & Melkonian, M. Centrin scaffold in Chlamydomonas reinhardtii revealed by immunoelectron microscopy. Eukaryot Cell 4, 1253–1263 (2005).

## References

1. Zhu, X., Wang, J., Li, S., Lechtreck, K. & Pan, J. IFT54 directly interacts with kinesin-II and IFT dynein to regulate anterograde intraflagellar transport. Embo j 40, e105781 (2021).

2. Girish, V. & Vijayalakshmi, A. Affordable image analysis using NIH Image/ImageJ. Indian J Cancer 41, 47 (2004).

3. Collins, T.J. ImageJ for microscopy. Biotechniques 43, S25–S30 (2007).

4. Mastronarde, D.N. Automated electron microscope tomography using robust prediction of specimen movements. J Struct Biol 152, 36–51 (2005).

5. Zheng, S.Q. et al. MotionCor2: anisotropic correction of beam-induced motion for improved cryo-electron microscopy. Nat Methods 14, 331–332 (2017).

6. Kremer, J.R., Mastronarde, D.N. & McIntosh, J.R. Computer visualization of three-dimensional image data using IMOD. J Struct Biol 116, 71–76 (1996).

7. Liu, Y.-T. et al. Isotropic Reconstruction of Electron Tomograms with Deep Learning. bioRxiv, 2021.2007.2017.452128 (2021).

8. Goddard, T.D. et al. UCSF ChimeraX: Meeting modern challenges in visualization and analysis. Protein Sci 27, 14–25 (2018).

